# Multiple routes to red-shifted chlorophyll *d*-based photosynthesis

**DOI:** 10.64898/2026.06.16.732656

**Authors:** Himanshu S. Mehra, Nikki Cecil M. Magdaong, David A. Flesher, Gaozhong Shen, Nikea J. Ulrich, Christian M. Brininger, Dariusz M. Niedzwiedzki, Scott R. Miller, Christopher J. Gisriel

## Abstract

Strains of the cyanobacterium *Acaryochloris marina* exhibit diverse far-red light-harvesting properties during chlorophyll *d*-based photosynthesis. Here, we show that differences in light absorption among *A. marina* strains arise exclusively from Photosystem I (PSI) and reflect variation in multiple low-energy chlorophyll states. Time-resolved fluorescence reveals different combinations of low-energy states among strains, generating a continuum of spectral phenotypes. Cryo-EM structures of PSI at ∼1.8 Å resolution reveal similar low-energy states arising from distinct pigment environments, demonstrating that red-shifted absorption is not governed by a single conserved motif. Phylogenetic analyses show that spectral tuning evolved through modular variation and reassortment of PSI components. These results indicate that distinct pigment configurations can converge on similar low-energy states, extending light harvesting near the energetic limit of oxygenic photosynthesis.

## Introduction

Oxygenic photosynthesis relies on pigment-protein complexes that harvest light and convert it into chemical energy (*1*). In most phototrophs, this process is mediated by chlorophyll *a*-based photosystems that primarily absorb visible light. However, the cyanobacterium *Acaryochloris marina* utilizes chlorophyll *d* as its primary photosynthetic pigment, enabling absorption of farred light (700–800 nm, **fig. S1**) (*2*). This adaptation allows *A. marina* to occupy ecological niches where longer-wavelength light predominates, e.g., shaded or spectrally filtered environments (*3*).

Spectroscopic analyses have revealed substantial light-harvesting diversity among *A. marina* strains (*4*–*8*). Notably, most previous studies have focused on strain MBIC11017 (hereafter MBIC), which represents a spectroscopic and genetic outlier relative to broader *A. marina* diversity. Across the genus, strains have been classified into short-, intermediate-, and long-wavelength spectral types, with low-energy absorption extending from ∼730 nm to ∼750–770 nm (6). These differences likely reflect adaptation to distinct far-red light environments (*6*), but the molecular mechanisms underlying the spectral diversity remain unclear. More broadly, this system highlights a central question in photosynthesis: how do pigment-protein environments tune chlorophyll excited-state energy landscapes near the red limit of oxygenic photosynthesis?

Photosystem I (PSI) is a pigment-protein complex that performs light-driven charge separation and is responsible for most low-energy absorption in oxygenic phototrophs. The structure of PSI from MBIC was previously determined (*9*) but lacked many peripheral features, limiting insight into energy tuning in chlorophyll *d*-based photosynthesis. More recently, a structure of PSI from the long-wavelength strain NIES-2412 (hereafter NIES) revealed substantial differences in a peripheral region centered on the PsaB subunit and the small transmembrane subunit PsaX (*5*). In NIES, PsaX is present, and PsaB adopts an architecture associated with closely spaced chlorophylls proposed to give rise to low-energy “red” states. In contrast, PSI from MBIC is thought to lack this arrangement and the associated red states. These observations have led to a model in which red-shifted absorption arises from a specific structural motif. However, the broader diversity of *A. marina* strains suggests that this model may be incomplete.

Here, we integrate biochemical characterization, steady-state and time-resolved spectroscopy, high-resolution cryo-electron microscopy (cryo-EM), and phylogenetic analyses to define the molecular basis of spectral diversity in *A. marina*. We show that differences in absorption arise solely from PSI and reflect variation in multiple low-energy chlorophyll states. By comparing PSI complexes across strains, we demonstrate that similar low-energy states can emerge from distinct structural configurations. These findings indicate that red-shifted photosynthesis in *A. marina* is achieved through multiple structural and evolutionary routes, revealing a flexible framework for tuning light harvesting near the red limit.

### Photosystem I defines spectral diversity in *Acaryochloris marina*

To investigate the basis of spectral diversity in *A. marina*, we analyzed representative strains spanning short-(MBIC), intermediate- (MU03 and MU05), and long-wavelength (FH6) spectral types. Whole-cell absorption and fluorescence at room temperature confirmed the expected spectral phenotypes of these strains (**Fig. 1A** and **fig. S2**). To identify the photosynthetic component(s) underlying these differences, we separated pigment-protein complexes using density gradient centrifugation (**Fig. 1B** and **fig. S3**). Each gradient resolved multiple chlorophyll-containing bands corresponding to distinct photosynthetic assemblies. Their absorption spectra revealed that, within a given strain, most components exhibited similar spectral profiles, whereas a single high-density component exhibited a pronounced red shift in the Q_y_ region (**fig. S3**).

**Fig. 1.**
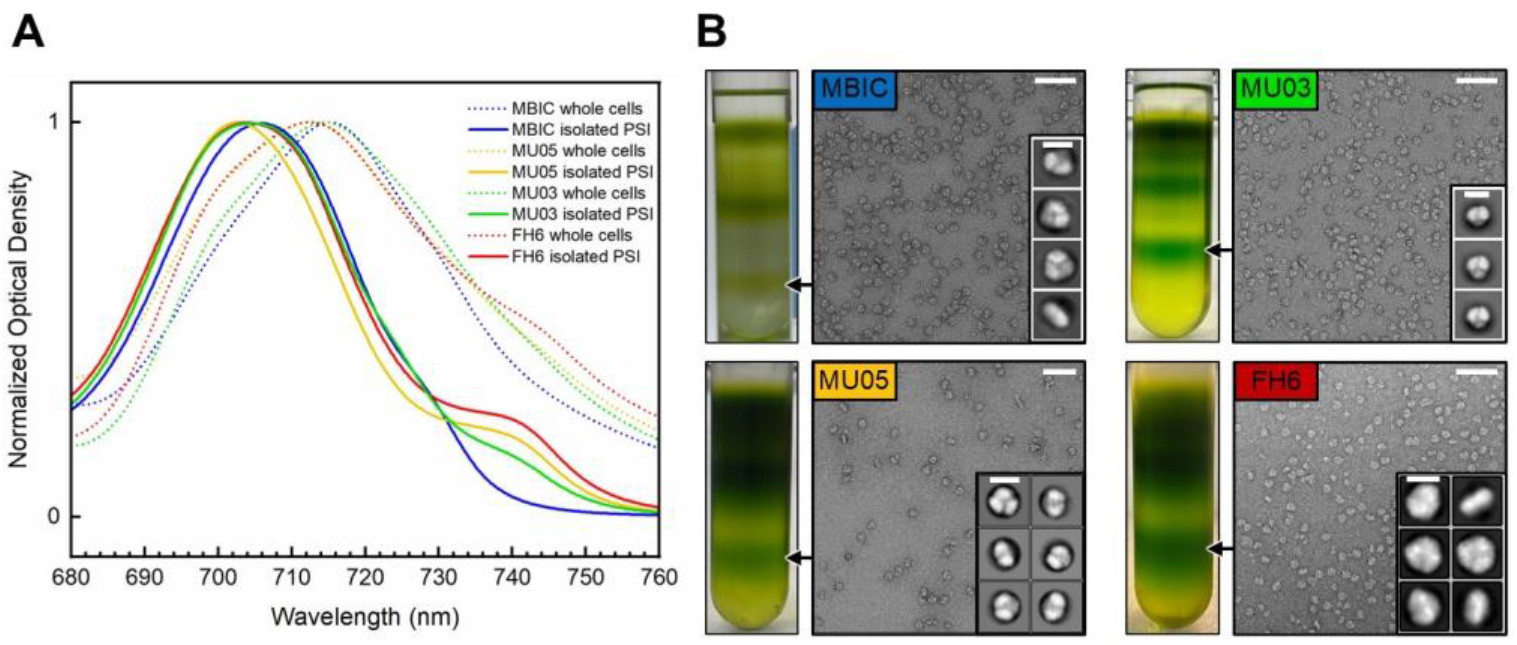
PSI underlies spectral diversity in *Acaryochloris marina*. (**A**) Normalized whole cell and PSI absorption spectra showing the chlorophyll *d* Q_y_ region. (**B**) Trehalose density gradients and TEM micrographs of the bottom bands. Scale bars associated with micrographs are 10 nm. 2D classes for each are also shown with a scale bar of 0.2 nm. Absorption spectra for each trehalose band are shown in **fig. S3**.

Transmission electron microscopy of this band after negative staining revealed homogeneous particles consistent with trimeric PSI (**Fig. 1B**). Inclusion of glutaraldehyde during gradient separation to stabilize potential PSI-antenna interactions did not reveal PSI particles associated with antenna subunits, and mass spectrometry showed that PSI subunits dominate these fractions, with only minor contributions from antenna proteins (**table S1**). The bottom bands on these gradients were therefore assigned solely as trimeric PSI complexes.

Comparison of isolated PSI spectra across strains revealed that all complexes share a Q_y_ maximum near ∼706 nm but differ in the extent of red-shifted absorption, with PSI from MU03, MU05, and FH6 exhibiting progressively enhanced red shoulders (**Fig. 1A**). These differences closely mirror the spectral phenotypes observed in whole cells, demonstrating that variation in PSI is sufficient to account for differences in absorption among *A. marina* strains. Consistent with this assignment, PSI subunits exhibit greater sequence divergence than those of PSII (**figs. S4**–**S5** and **tables S2**– **S3**).

### Steady-state spectroscopy reveals strain-dependent low-energy states

Steady-state spectroscopy of isolated PSI complexes revealed distinct low-energy absorption features that vary among *A. marina* strains. 77 K absorption measurements enabled resolution of multiple bands in the chlorophyll *d* Q_y_ region, corresponding to low-energy chlorophyll populations (**Fig. 2A**). Second-derivative analysis identified conserved components near ∼700– 711 nm in all strains, along with additional strain-dependent features at longer wavelengths. MBIC exhibits a single low-energy band near ∼730 nm. MU03 contains this feature together with additional lower-energy bands near ∼740 and ∼745 nm, whereas MU05 and FH6 lack the ∼730 nm component and instead resemble only the lower-energy portion of the MU03 spectrum.

**Fig. 2.**
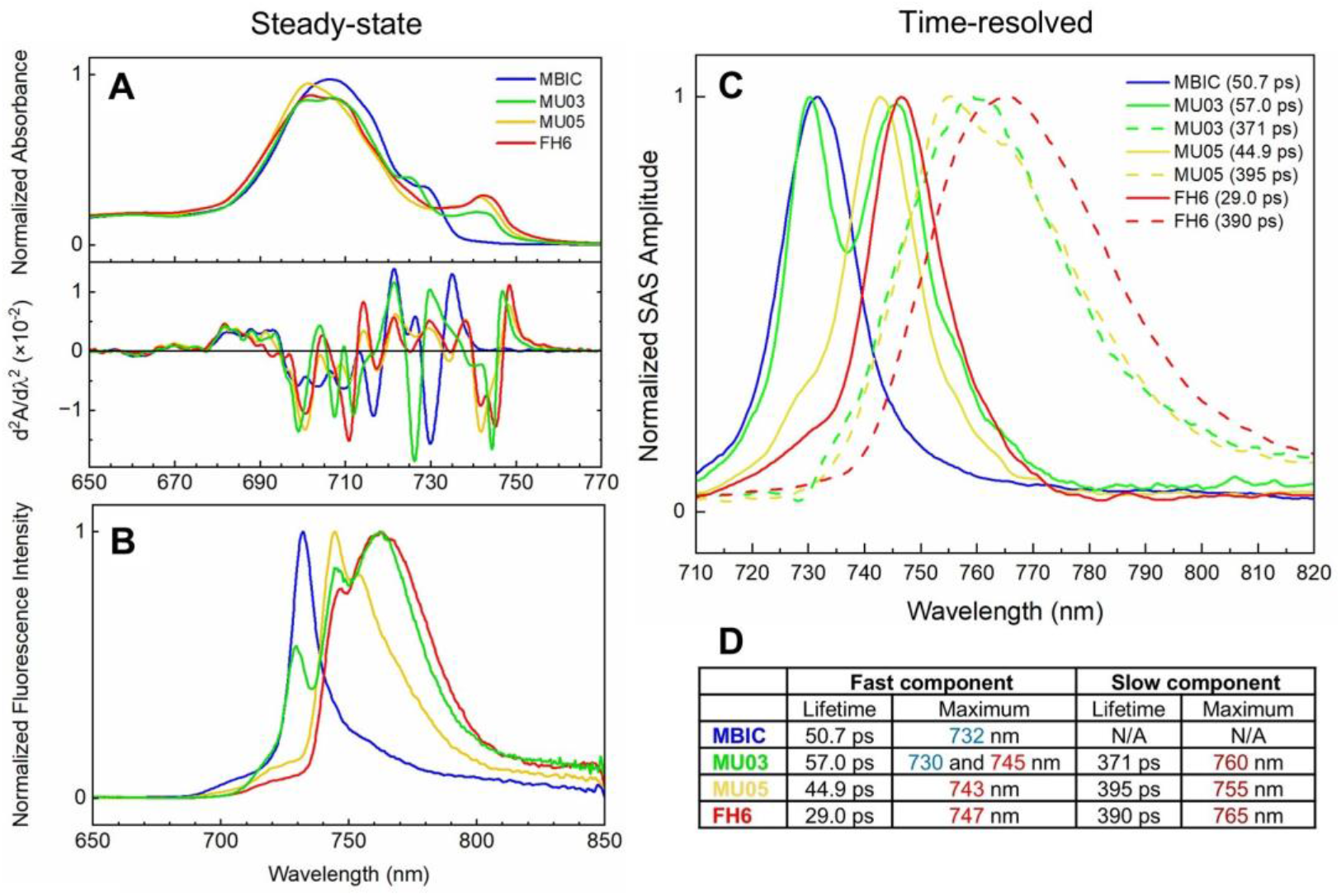
Spectroscopic characterization of *A. marina* PSI. (**A**) 77 K absorption in the chlorophyll *d* Q_y_ region and the corresponding second derivative analysis. (**B**) 77 K fluorescence emission spectra (excitation wavelength = 490 nm). (**C**) Global analysis of 77 K TRF of spectral variants of PSI. Species associated spectra (SAS) normalized to unity. Components with ns-scale kinetic lifetimes corresponding to free chlorophylls are excluded for clarity. (**D**) Tabulated data from panel C.

Fluorescence emission spectra mirror these energetic differences (**Fig. 2B**). At 77 K, MBIC PSI emits predominantly near ∼730 nm, whereas MU03 exhibits this feature together with additional lower-energy emission components. MU05 and FH6 lacked the ∼730 nm emission and instead displayed only the lower-energy states shared with MU03. These observations indicate that MBIC contains a single dominant terminal emitter, whereas MU03, MU05, and FH6 contain multiple low-energy fluorescence states.

The previous work on NIES PSI assigned two red states fluorescing at 745 and 765 nm (*5*), whose PSI subunit sequences are identical to those of strain FH6 in our analysis. The spectral features observed here are broadly consistent with this framework and support the presence of multiple lower-energy states in intermediate- and long-wavelength strains. The relative contributions of these states vary among strains, indicating that the low-energy landscape of *A. marina* PSI is modulated in a strain-dependent manner.

### Time-resolved fluorescence supports diversity in low-energy states among strains

To determine how low-energy states contribute to excitation energy dynamics in PSI, we performed 77 K time-resolved fluorescence (TRF) measurements on isolated complexes from each strain (**Fig. 2C** and **text S1**). Global analysis revealed kinetic components that report on excitation energy flow through distinct low-energy states (**fig. S6**).

All PSI complexes exhibit a fast fluorescence component with 30–60 ps lifetime centered near 730 nm and/or 745 nm depending on the strain (**Fig. 2C**). In MBIC, this component was dominated by an ∼730 nm band, consistent with the single dominant low-energy emitter observed in the steady-state fluorescence spectrum (**Fig. 2B**). In contrast, MU05 and FH6 display fast components shifted toward ∼745 nm, indicating that the dominant low-energy fluorescent state in these strains is red-shifted relative to MBIC. Interestingly, MU03 exhibits both the ∼730 nm and ∼745 nm fast components, strongly implying that these states arise from separate sites rather than from a single site that is differently tuned among strains.

In addition to these fast components, PSI from MU03, MU05, and FH6 (but not MBIC) exhibit a slower fluorescence component (300–400 ps lifetime) with emission maxima between 755–765 nm (**Fig. 2C**). The slower decay and red-shifted emission are consistent with a population of a very low-energy state after equilibration. Together, these data support a model in which PSI contains multiple terminal low-energy states whose relative contributions vary among strains. The absence of a slower, more red-shifted component in MBIC indicates that, at 77 K, excitation remains localized to a site with higher-energy than the P_740_ trap. The substantially larger Stokes shift of the slow emitter relative to the fast emitters (**table S4**) further suggests a more flexible structural environment for the lowest-energy state.

### High-resolution structures reveal candidate origins of low-energy chlorophyll states

To interpret the spectroscopic properties of PSI, we determined cryo-EM structures of PSI from strains MU03 and MU05 at resolutions of 1.81 and 1.77 Å. Data collection, processing, model building, cofactor assignments, and model statistics are provided in **text S2, figs. S7**–**S12**, and **tables S5**–**S6**. The high resolution of these structures enabled detailed examination of pigment environments and evaluation of previous hypotheses regarding cofactor composition, while providing a structural framework for the analyses described below.

The major structural differences between MU03 and MU05 PSI are concentrated in the peripheral region of PsaB near where PsaX binds. Here, the arrangement of chlorophylls differs substantially between the two structures (**Fig. 3A**). To help identify possible low energy chlorophyll sites that correspond to spectroscopic features, we (i) identified structural differences that could influence site energies, and (ii) calculated approximate Förster energy transfer rates between chlorophylls based on ring geometries (**Fig. 43** and **data S1**) (*10*) because strong pigment coupling is associated with red-shifted states.

**Fig. 3.**
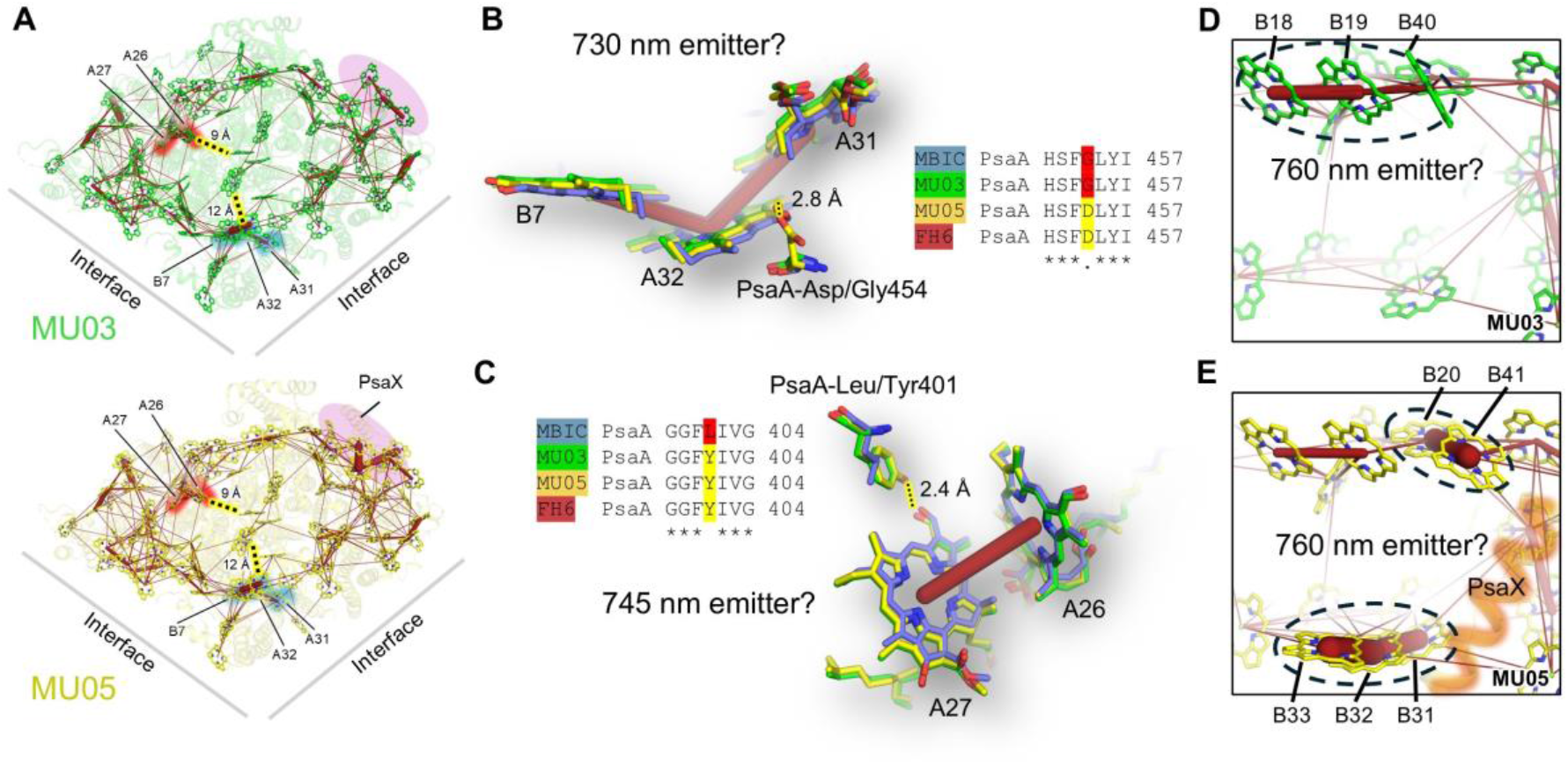
Structural analysis of red sites in PSI. (**A**) Monomeric structures of MU03 and MU05 PSI. Lines between chlorophylls denote faster (thick) to slower (thin) calculated energy transfer rate based on the Förster rate analysis. Chlorophylls implicated in emitting sites are labeled. The region highlighted in purple corresponds to the major structural difference between MU03 and MU05 PSI. (**B**) Close-up on chlorophylls B7, A32, and A31 from superimposed structures of MBIC, MU03, and MU05 PSI proposed to give rise to the 730 nm fluorescence emission. The hydrogen bond interaction proposed to tune the energy of this site is shown with its corresponding partial sequence alignment. Lines between chlorophylls denote calculated energy transfer rates as in panel A. (**C**) Close-up on chlorophylls A26 and A27 proposed to give rise to 745 nm fluorescence emission. The hydrogen bond interaction proposed to tune the energy of this site is shown with its corresponding partial sequence alignment. (**D**) and (**E**) Peripheral regions of PsaB corresponding to those highlighted in purple in panel B. Chlorophyll sites proposed to be responsible for fluorescence emission at ∼760 nm are circled.

Due to their fast fluorescence lifetime (**Fig. 2C**) and small Stokes shift (**table S4**), we searched the structural data for potential fluorescence emitters near the electron transfer chain. A candidate for the ∼730 nm TRF component is the B7/A32/A31 chlorophyll trimer (chlorophyll site names are as in Jordan et al. (*11*)), which is well-coupled and located near the electron transfer chain (**Fig. 3A** and **3B**). Strains that contain the ∼730 nm component (MBIC and MU03) have PsaA-Gly454 near the A32 C3 formyl group, whereas MU05 and FH6 contain Asp. Notably, the sister taxon *Acaryochloris thomasi* also contains Gly at this position, suggesting that this is the ancestral state. The Gly-to-Asp substitution at this position in MU05 and FH6 PSI could alter the site energy or coupling of the trimer, providing a plausible structural basis for the absence of the ∼730 nm component in MU05 and FH6. It is also worth noting that the B7/A31/A32 site has been proposed as a low energy site in chlorophyll *a*-containing PSI (*12, 13*).

A candidate for the ∼745 nm TRF component is the A27/A26 chlorophyll pair, which is well coupled and lies ∼9 Å from P_740_ in the electron transfer chain (**Fig. 3A** and **3C**). In MU03, MU05, and FH6, PsaA-Tyr401 donates a hydrogen bond to the C3 formyl group of A27, an interaction absent in MBIC. Interestingly, *A. thomasi* contains Ile at the corresponding position suggesting that the ancestral state may have resembled the higher-energy MBIC configuration. In any case, because hydrogen bonding to the formyl moiety is expected to lower the site energy, the strain distribution of this interaction supports assignment of the A27/A26 pair to the 745 nm TRF component.

Assignment of the lowest-energy TRF component (∼760 nm) is more challenging. This component is observed in MU03, MU05, and FH6 and exhibits a large Stokes shift and long lifetime, consistent with emission from a peripheral, low-energy state. Structural differences among strains are concentrated in the stromal-side region of PsaB near the PsaX binding site (**Fig. 3D** and **3E**). In MU05 and NIES PSI structures, this region contains the B20/B41 dimer and B31/B32/B33 trimer, which have been proposed to give rise to low-energy “red” states and appear strongly coupled in our Förster energy transfer analysis. In contrast, MU03 lacks PsaX and these chlorophyll clusters, instead exhibiting a distinct trimer, B18/B19/B40, in a similar location as the B20/B41 dimer.

Despite these structural differences, MU03 exhibits an ∼760 nm TRF component with wavelength and lifetime comparable to MU05 and FH6. This suggests that the lowest-energy state does not require a specific conserved chlorophyll cluster but instead arises from alternative pigment-protein configurations within this region. By analogy to the proposed assignment of B20/B41 in NIES, the B18/B19/B40 trimer in MU03 provides a plausible candidate for the ∼760 nm component, indicating that different structural arrangements can produce similar low-energy states.

Additional support for localization of the ∼760 nm component near this region comes from steady-state spectroscopic comparison of PSI trimers and monomers (**fig. S13**). The low-energy features in absorption and fluorescence are substantially decreased in monomers, consistent with disruption of a low-energy state near the monomer-monomer interface. Although the structures of the monomeric complexes are unknown, monomerization could destabilize the adjacent PsaB region implicated in formation of the lowest-energy states. Interpretation of this region is nevertheless complicated by partial disorder and reduced local resolution (**text S2**), particularly in MBIC and MU03. This is consistent with the structural flexibility expected for peripheral low-energy states. Uncertainty in chlorophyll orientation therefore limits detailed interpretation of coupling interactions, and sequence divergence relative to chlorophyll *a*-containing PSI constrains inference by analogy. Nevertheless, the sensitivity of the ∼760 nm component to monomerization, together with its consistent appearance across structurally distinct PSI complexes, supports a model in which the lowest-energy state emerges from a conserved spatial region rather than a specific chlorophyll arrangement.

### Phylogenetics reveals modular evolution of PSI spectral tuning

Because the PsaX-associated region strongly influences the lowest-energy fluorescence component and likely contributes substantially to the whole-cell fluorescence phenotype, we sought to place the structural and spectroscopic differences among PSI complexes in an evolutionary context. Reconstruction of PsaX relationships among diverse *A. marina* strains revealed six major allele families (**Fig. 4A**). The distribution of these alleles across the species phylogeny (**fig. S14**) indicates a history of genetic exchange and reassortment among divergent genomic backgrounds. Among PsaX-containing strains, allele identity is associated with the extent of red-shifting: strongly red-shifted strains such as FH6 and NIES belong to distinct PsaX allele groups compared to intermediate strains such as MU05. The relatively low sequence identity between PsaX in MU05 and in FH6 or NIES (50%, **table S3**) suggests that divergence at this locus contributes to tuning of low-energy states within this region of PSI, consistent with our assignment of the lowest-energy TRF component to the adjacent PsaB/PsaX-associated region.

**Fig. 4.**
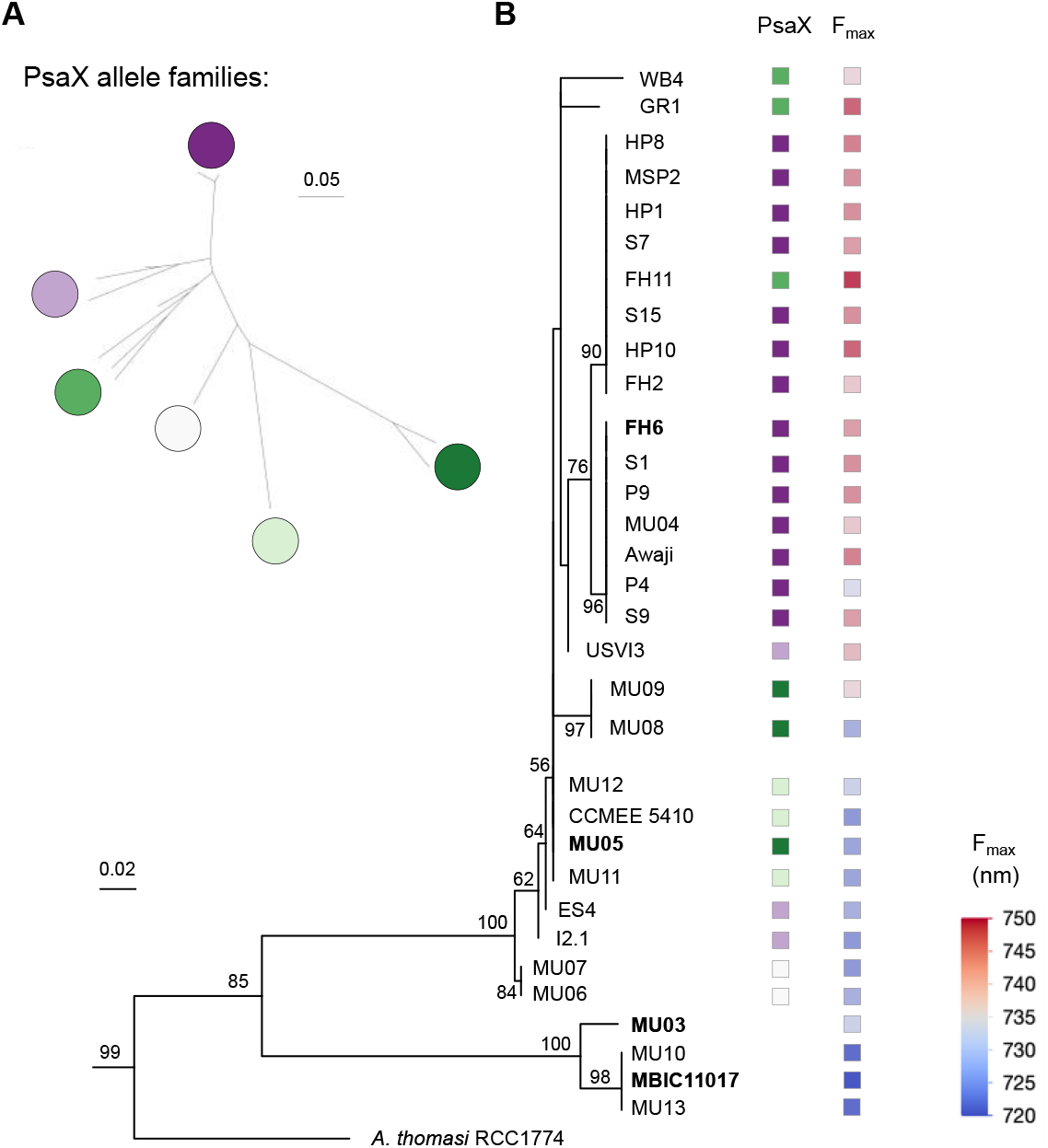
Evolutionary diversification of low-energy states in *A. marina* PSI. (**A**) Relationships among *A. marina* PsaX alleles. (**B**) Maximum likelihood tree of the PsaB region spanning residues 250–530, which includes the structural elements implicated in potential low-energy chlorophyll environments. Strains characterized in this study are indicated in bold. PsaX allele families are mapped onto the tree (color coding as in panel A) along with room temperature fluorescence maxima, which ranges from higher (blue) to lower (red) energy level. For both analyses, branch lengths are in units of expected number of amino acid substitutions per site.

To determine whether variation in the PSI core also contributes to spectral diversity, we analyzed the evolutionary history of full-length PsaA and PsaB (**fig. S15**), as well as the PsaB region encompassing residues 250–530 (**Fig. 4B**) that contains the structural elements implicated in the lowest-energy chlorophyll environments (**Fig. 3D–E**). Like PsaX, both PsaA and PsaB exhibit signatures of genetic exchange, resulting in topological differences compared with the genome-wide phylogeny (**figs. S14** and **S15**). Distantly related PsaX-lacking strains (MU03 and the blue-shifted strains) form a distinct PsaB lineage that is highly divergent from PsaX-containing strains (**Fig. 4B**); for the latter, more red-shifted strains tend to possess more derived PsaB alleles. Notably, strains with similar PsaX alleles but different PsaB alleles can exhibit substantially divergent fluorescence maxima. Consequently, differences in spectral tuning reflect interactions between PsaX variation and the surrounding PSI core structural background.

The two PsaB regions that interact with PsaX also exhibit contrasting patterns of diversification. Since diverging from the *A. marina* common ancestor, the PsaX-containing strains have evolved faster in the region encompassing residues 250–350, whereas the PsaX-lacking lineage has evolved faster at residues 450–530 (**figs. S16** and **S17**). Thus, the structural modules associated with the lowest-energy chlorophyll environments appear to have followed distinct evolutionary trajectories during *A. marina* diversification. These observations suggest that different lineages modified different components of the same region to generate low-energy states, providing an evolutionary explanation for the multiple structural solutions identified in the cryo-EM analyses.

Taken together, these data support a model in which red-shifted spectral diversity in *A. marina* arises through modular evolution of PSI. A PsaX-independent pathway, represented by MU03, enables formation of low-energy states through tuning of the PSI core scaffold. A second pathway involves diversification and reassortment of PsaX together with adjacent PSI-core features, generating distinct combinations of structural elements that modulate the extent of red-shifting. Thus, similar low-energy states can convergently emerge through multiple evolutionary routes acting on the same region of PSI.

## Discussion

This work provides a framework for understanding how low-energy states evolve in photosystems operating near the red limit of oxygenic photosynthesis. Whereas previous work emphasized specific pigment arrangements as the origin of red-shifted states, our results show that similar spectral features can emerge from distinct structural configurations, indicating a flexible relationship between structure and energetics. This conclusion is consistent with decades of work on chlorophyll *a*-containing PSI, where multiple candidate red sites have been proposed (*11*–*19*). Notably, introduction of the PsaB loop required for B33 binding is sufficient to generate a red-shifted state in PSI from *Synechocystis* sp. PCC 6803, even in the absence of PsaX (*12*). Our findings extend this principle to chlorophyll *d*-based PSI, demonstrating that divergent pigment-protein environments can converge on similar energetic outcomes.

Within *A. marina*, the observed diversity likely reflects a history of modular evolution and horizontal exchange of PSI components. Variation in PsaX correlates with differences in spectral properties among PsaX-containing strains, yet the presence of strongly red-shifted states in MU03 demonstrates that PsaX is not required for their formation. Instead, reassortment of divergent PsaX alleles across distinct PSI core backgrounds during *A. marina* diversification may have generated novel combinations of structural features that produce alternative low-energy states. The association between spectral phenotype and variation in the adjacent PSI core further suggests that low-energy states evolve through modular combinations of PSI structural elements rather than changes in PsaX alone.

The phylogenetic analyses further indicate that the structural modules associated with the lowest-energy chlorophyll environments followed distinct evolutionary trajectories. The PsaB regions implicated in low-energy states exhibit contrasting patterns of diversification in PsaX-containing and PsaX-lacking lineages, suggesting that different lineages modified different components of the same structural region. This provides an evolutionary explanation for the multiple structural solutions identified in the cryo-EM analyses, in which similar energetic outcomes arise through different combinations of PSI features.

The absence of PsaX in the distantly related sister taxon *A. thomasi* suggests that the ancestral *A. marina* PSI may also have lacked PsaX, with PsaX-containing architectures arising later in some lineages. At the same time, the shared structural features between MU03 and short-wavelength strains imply genetic exchange involving PSI gene cluster components, although the directionality of these events remains unclear. One possibility is that distinct PSI gene cluster configurations, including MBIC/MU03-like and MU05/FH6/NIES-like states, were exchanged during diversification of the lineage, generating alternative structural templates for low-energy states. Distinguishing among these possibilities will require broader genomic sampling and reconstruction of PSI gene cluster evolution.

Our results also suggest that the traditional classification of *A. marina* strains into short-, intermediate-, and long-wavelength spectral types oversimplifies the underlying diversity of PSI energetics. Rather than discrete spectral classes, the spectroscopic, structural, and phylogenetic data support a continuum of low-energy configurations generated through different combinations of PSI features. MU03 illustrates this particularly well, combining low-energy states associated with both MBIC-like and FH6/NIES-like PSI architectures. Spectral diversity in *A. marina* is therefore better viewed as a multidimensional energetic landscape than as a small number of discrete phenotypic classes.

Beyond the structural and evolutionary mechanisms discussed above, the use of chlorophyll *d* introduces an additional level of spectral tuning through its C3 formyl group, which can modulate site energies through hydrogen bonding and electrostatic interactions, as illustrated by the B7/A32/A31 and A27/A26 sites identified here. In addition, the use of pheophytin *a* in the electron transfer chain implies altered charge-separation energetics relative to canonical PSI (*20*), potentially influencing excitation-energy transfer to the reaction center.

Several limitations of the present study should be considered. While the combined structural, spectroscopic, and phylogenetic analyses strongly implicate the PsaB/PsaX-associated region in formation of the lowest-energy state, assignment of individual chlorophylls to specific spectroscopic components remains provisional. Future studies employing targeted mutagenesis, time-resolved infrared spectroscopy, and excited-state calculations should help distinguish among the candidate sites proposed here and further define the basis of red-shifted light harvesting in chlorophyll *d*-based PSI.

Taken together, our results suggest that low-energy states in PSI are not defined by a single conserved structural motif but instead arise from a spatially and energetically flexible pigment-protein environment. This flexibility allows multiple structural solutions to converge on similar outcomes, enabling adaptation to diverse light environments near fundamental energetic limits. Yet the repeated involvement of the same PsaB/PsaX-associated region indicates that these solutions emerge from a constrained structural framework imposed by the PSI core architecture.

## Acknowledgments

We thank the Cryo-EM Research Center (CEMRC) in the Department of Biochemistry, the 3D Cell Electron Microscopy Core Facility, and the Biotechnology Center Mass Spectrometry Core Facility at the University of Wisconsin-Madison for instrument use and technical assistance. We thank the Laboratory for BioMolecular Structure (LBMS) at Brookhaven National Laboratory for use of the Krios facility. The LBMS is supported by the DOE Office of Biological and Environmental Research (KP160711). We thank the Ultrafast Laser Facility (ULF) in the Department of Biology at Washington University in St Louis. The ULF is supported by U.S. Department of Energy, Office of Science, Office of Basic Energy Sciences grant DE-FG02-99ER20350. We thank Yuval Mazor for assistance with Förster energy transfer calculations.

## Author contributions

Conceptualization: SRM, CJG

Methodology: HSM, NCMM, DAF, GS, NJU, CMB, DMN, SRM, CJG

Investigation: HSM, NCMM, GS, NJU, CMB, DMN, SRM, CJG

Writing – original draft: HSM, CJG

Writing – review & editing: HSM, NCMM, DAF, GS, NJU, CMB, DMN, SRM, CJG

Funding acquisition: DMN, SRM, CJG

Resources: DMN, SRM, CJG

Supervision: SRM, CJG

## Data, code, and materials availability

Cryo-EM maps and coordinates have been deposited in the Protein Data Bank (PDB) and Electron Microscopy Data Bank (EMDB) under accession numbers 35ZQ/EMD-77301 and 36AI/EMD-77326 for MU03 and MU05 PSI, respectively. Additional spectroscopic data supporting the findings of this study are provided in the supplementary materials. Additional data are available from the corresponding authors upon request.

## Supplementary Materials

Materials and Methods

Supplementary Text 1–3

Figs. S1 to S17

Tables S1 to S6

References 5, 6, 9, 11, 21–43

Data S1

