## Supplemental Information for "Multiple routes to red-shifted chlorophyll *d*-based photosynthesis"

**The file includes:**

Materials and Methods

Supplementary Text S1 to S3

Figs. S1 to S17

Tables S1 to S6

References 5, 6, 9, 11, 21–43

**Other Supplementary Materials for this manuscript include the following:**

Data S1

### Materials and Methods

Cell growth and PSI isolation

*A. marina* strains MU03, MU05, and FH6 were grown in Instant Ocean BG-11 medium with components as described in Ulrich et al. (*6*) and MBIC was grown in modified ASNIII medium as described in Gan et al. (*21*). The cultures were maintained by slowly sparging air and gently stirring at 25 °C under 10–15 μmol photons $m^{-2}$ $s^{-1}$ of cool white, fluorescent light.

For PSI isolation, cell pellets were harvested and washed once in 50 mM MES pH 6.5 with 20 mM $CaCl_2$ and 10 mM $MgCl_2$. Cells were pelleted again and resuspended in the MES buffer with a cell-to-buffer ratio of 1 g cells (wet weight) in 10 mL of MES buffer. Cells were lysed by using bead beating and a continuous flow cell disruptor (Constant Systems). Unbroken cells were removed via centrifugation at 4,284×*g*. Thylakoid membranes were pelleted by ultracentrifugation at 126,000×*g* for 30 min and were resuspended in 50 mM MES pH 6.5 with 20 mM $CaCl_2$, 10 mM $MgCl_2$, and 5% trehalose. Membranes were then solubilized in the dark at 4 °C for 30 min by adding 1% (w/v) *n*-dodecyl β-d-maltoside (β-DDM). The insoluble debris was removed by centrifugation at 10,967×*g*. The solubilized membranes were loaded onto 8 to 22% trehalose gradients, which were prepared with 50 mM MES pH 6.5 with 20 mM $CaCl_2$, 10 mM $MgCl_2$, and 0.03% (w/v) β-DDM with or without 0.1% (v/v) glutaraldehyde. Gradients were ultracentrifuged for 17 h at 108,000×*g* at 4 °C. The lowest green-colored bands, which contained PSI trimers, were collected and concentrated using an Amicon 50 kDa centrifugal filter (EMD Millipore). The concentrated PSI sample was then loaded onto a second 8 to 22% trehalose gradient containing 0.03% β-DDM and subjected to a second round of ultracentrifugation. The purified trimeric PSI band was collected from the gradients.

Negative staining and transmission electron microscopy

4-μL of protein at ~1–2 μg chlorophyll/mL was pipetted onto either a glow-discharged (25 mA, 30 s) Cu Carbon 400 mesh (Ted Pella) or a Cu Formvar/Carbon 400 mesh (Electron Microscopy Sciences) microscopy grid and incubated for 1 min at room temperature. The liquid was wicked away using filter paper, and the grid was then washed two times in 20 μL drops of 50 mM MES pH 6.5 with 20 mM $CaCl_2$, 10 mM $MgCl_2$, and 0.03% β-DDM. The same procedure was repeated two times with 20 μL drops of 2% uranyl acetate solution, and the grid was then incubated on a final 20 μL stain drop for 1 min. The liquid was wicked away, and the grid was blotted to complete dryness. The prepared grids were stored in a vacuum chamber before imaging. The sample grids were imaged using either a 200 kV Talos F200C or a 120 kV Talos L120C, and micrographs were collected for data processing. The micrographs were processed using either RELION 5.0 (*22*) or CryoSPARC v4.7.1 (*23*).

Steady-state spectroscopy

Steady-state absorption spectra were recorded using Shimadzu UV-Vis spectrophotometers (models 1900i and UV-Vis 1800). Fluorescence emission spectra were measured using a HORIBA Fluorolog-QM spectrofluorometer equipped with double monochromators and a photomultiplier tube detector. For cryogenic measurements, PSI samples were mixed with glycerol to a final concentration of 60% (v/v) and cooled in a liquid nitrogen vapor cryostat (Janis VNF-100), where they formed transparent glasses at low temperature. Spectra at 77 K were recorded using appropriate cryogenic cuvettes.

Time-resolved fluorescence spectroscopy

Fluorescence decay maps were measured with a streak camera system (Hamamatsu, Japan), consisting of a universal C5680 streak camera and spectrograph from Bruker described in detail previously (*24*). It was coupled to the ultrafast-laser system based on an ultrafast Ti:Sapphire laser (Mai-Tai, Spectra-Physics, USA), an ultrafast optical parametric oscillator (Inspire100, Spectra-Physics, USA) and a pulse selector (Model 3980, Spectra-Physics, USA), used to lower the frequency of the excitation beam to 8 MHz. The excitation beam was set to 410 nm (exciting B bands of chlorophyll *d*), focused on a circular spot of <1 mm diameter, depolarized prior to the sample using an achromatic depolarizer (DPU-25, Thorlabs, USA), and set to low photon density of ~$10^{10}$ photons/$cm^2$ per pulse that warranted annihilation-free excitation migration in large pigment arrays as of those in PSI proteins. To avoid light scattering effects, a 610 nm long-pass filter was placed at the spectrograph entrance.

<u>Global analysis of time-resolved data</u>

The TRF datasets were globally fitted using a parallel decay model according to:

$$Fl(t,\lambda) = \sum_{i=1}^{n} DAS_i(\lambda)\, C_i(t)$$

where $DAS$ denotes decay-associated spectra and $C_i(t)$ represents the time-dependent concentration of each component, defined as $C_i(t) = IRF \otimes e^{-t/\tau_i}$, where $IRF$ is the instrument response function, $\tau_i$ is the lifetime of component $i$, and $\otimes$ denotes convolution. This model assumes simultaneous excitation and independent decay of all excited components.

When the resulting spectral components corresponded to physically meaningful fluorescence spectra, DAS components were interpreted as species-associated spectra (SAS). Steady-state fluorescence spectra were reconstructed by integrating the time-resolved signal over the fluorescence decay:

$$F(\lambda) = \sum_{i=1}^{n} SAS_i(\lambda) \int_0^{\infty} C_i\,(t)\, dt$$

Global analysis of TRF datasets was performed using CarpetView software (Light Conversion, Lithuania). The instrument response function was approximated by a Gaussian with a full width at half maximum (FWHM) of ~65 ps at 77 K. Data visualization was performed using Origin (OriginLab Corp., USA).

<u>Pigment extraction and HPLC analysis</u>

Pigment analyses were carried out under dim light and with samples kept on ice. All solvents used were HPLC-grade. Briefly, PSI samples were extracted with ~1 mL of 7:2 (v/v acetone/methanol). After centrifugation at 11,000×*g* for 2 min, the supernatant was collected and dried under $N_2$ gas. The sample was then resuspended in methanol, filtered (0.20 µm), and injected into an Agilent 1100 HPLC with a Zorbax Eclipse XDB-C18 column. The mobile phase was set to 99% methanol and 1% ethyl acetate for the first 10 min, then switched to 60% methanol and 40% ethyl acetate for the next 15 min and maintained until the end of the run (35 min total). Pigment quantitation was carried out in Chemstation for LC 3D Systems (Rev. B.04.03[16]) and Origin Pro 2025 by converting the peak area of each component detected at their absorption maximum to molar concentrations using their molar extinction coefficients at the corresponding wavelengths.

Mass spectrometry analysis

The purified protein complexes from the trehalose density gradients were concentrated in an Amicon 50 kDa centrifugal filter (EMD Millipore). An ~20 µg aliquot of the concentrated protein was used for 'in-liquid' Trypsin/LysC digestion and peptides were analyzed by nanoLC-MS/MS using the Agilent 1100 nanoflow system (Agilent) connected to hybrid linear ion trap-orbitrap mass spectrometer (LTQ-Orbitrap Elite™, Thermo Fisher Scientific) equipped with an EASY-Spray™ electrospray source.

Cryo-EM sample preparation, data collection, processing, and model building

Purified PSI complexes were applied to glow-discharged (25 mA for 30 s) Quantifoil R 1.2/1.3 Cu 300-mesh grids for MU03-PSI and Quantifoil R 2/1 Au 300-mesh grids for MU05 PSI. Grids were vitrified using a Vitrobot Mark IV (Thermo Fisher Scientific) operated at 4 °C and 100% humidity. Grids were blotted for 3 s and plunge-frozen in liquid ethane before transferring to liquid nitrogen. Initial grid screening was performed on an Arctica transmission electron microscope (Thermo Fisher Scientific, FEI) to assess particle distribution and ice quality.

High-resolution cryo-EM datasets were collected on a Titan Krios G3 transmission electron microscope (Thermo Fisher Scientific, FEI) operated at 300 kV and equipped with a Gatan K3 direct electron detector and energy filter (20 eV slit width for MU03 PSI and 15 eV slit width for MU05 PSI). Movies were recorded in super-resolution mode at a pixel size of 0.417 Å for MU03 PSI) and 0.413 Å for MU05 PSI, with a defocus range of approximately −0.7 to −1.9 µm in both cases. For both data sets, each exposure consisted of 50 frames with a total electron dose of 50.0 $e^{-}$ $Å^{-2}$. Automated data acquisition was performed using EPU. 15,366 micrograph movies were collected for MU03 PSI and 17,368 micrograph movies were collected for MU05 PSI.

Cryo-EM data processing was carried out using CryoSPARC v4.7.1, v5.0.2, and v5.0.4 (*23*). Data processing workflows and resolution estimations are provided in **fig. S7** and **fig. S8**. Micrograph movies were corrected and aligned using Patch motion correction. The contrast transfer function (CTF) was estimated using CTFFIND4 (*25*). The micrographs were then curated manually and denoised. The initial set of particles was picked using the blob-picker and filtered using the micrograph junk detector. The filtered particles were then inspected using the inspect particle picks job. The particles were then extracted using a box size of 420. Iterations of 2D classifications, ab initio reconstruction, and 3D refinements were performed to generate a good initial model, and the corresponding best 2D classes were used as templates for template picker. The same approach of particle filtering using the micrograph junk detector, inspect particle picks, multiple iterations of 2D classification, ab initio reconstruction, and 3D refinement was used to pick the best particles. The best particles from the blob picker and template picker were combined, and duplicate particles within 100 Å of each other were removed. Another round of particle filtering was performed, and high-quality reference volumes were generated using 3D refinements. The best particles were then corrected for per-particle motion, and empirical dose weights were estimated using the high-quality reference volumes by employing reference-based motion correction (*26*, *27*). Initially, a box size of 420 was selected for Fourier-cropping particles, but subsequent 3D refinements showed that the Fourier Shell Correlation (FSC) curves approached the Nyquist limit. A box size of 500 was then selected, and reference-based motion correction was performed again. 3D refinements after the reference-based motion correction (with Fourier crop box size 420) were examined using

ChimeraX (*28*) and WinCoot (*29*) and used as high-quality initial reference volumes for 3D refinements after the reference-based motion correction (with Fourier crop box size 500). The resulting non-uniform refinements yielded 3D reconstructions at 2.07 Å with 345,167 particles for MU03-PSI and at 1.97 Å with 592,088 particles for MU05-PSI. These particle sets were then symmetry expanded using $C_3$ symmetry. PSI monomer masks were generated in ChimeraX. Local refinements in $C_1$ symmetry were then performed using the best non-uniform refinement, symmetry-expanded particles, and the monomer mask. This resulted in PSI monomer 3D reconstructions at 1.93 Å for MU03 PSI and 1.83 Å for MU05 PSI. Using monomer masks, FSC validation was performed, yielding final resolution estimates of 1.81 Å for MU03 PSI and 1.77 Å for MU05 PSI. All map resolutions were calculated using the Gold-standard FSC of 0.143. Local resolution estimate and local resolution filtering were then performed, the latter of which was considered the final map for modeling.

For better resolving the peripheral region of PsaB, we generated peripheral masks for these regions in ChimeraX and ran local refinement jobs. The local refinements in the peripheral PsaB region for MU03 PSI yielded a 2.06 Å map, and in the peripheral PsaB/PsaX region for MU05 PSI yielded a 2.18 Å map. Local resolution estimation and local filtering were then performed for the peripheral regions.

Models were generated using the local filtered maps. The local filtered maps containing the full PSI monomers were selected as the main maps for Protein Data Bank/Electron Microscopy Data Bank deposition. The following maps were additionally uploaded: local refinement maps (sharpened and unsharpened), the half maps from the subsequent FSC validation jobs, the unsharpened focused map on the peripheral region from local refinement, and the local filtered focused map.

Initial models were generated based on homology modeling with PDB 7COY (*9*). Models were assembled and fit into the cryo-EM maps using PyMOL (*30*) and ChimeraX (*28*). Manual refinement was performed using Coot (*29*). Automated refinement was performed using real space refinement (*31*) in the Phenix software suite (*32*).

Analysis of tetrapyrrole type in the electron transfer chains of MU03 and MU05 PSI

To investigate the tetrapyrrole type in the $P_B$, $A_{-1A}$, and $A_{-1B}$ positions of the electron transfer chain from the two structures, a probe-atom sampling approach was developed based on the cone scan method reported previously (*33*, *34*). For each tetrapyrrole analyzed, the macrocycle was extracted from the atomic model and a series of probe atoms were placed in a cone shape extending outward from the C3 substituent attachment site (**fig. S10A**). Probe positions were generated using the vector from atom C3B to atom CAB, corresponding to the direction of the C3 substituent. The cone origin was placed at CAB and extended outward along this axis.

The cone geometry was chosen to approximate the expected orientations of either a formyl group (chlorophyll *d*) or a vinyl group (chlorophyll *a* or pheophytin *a*). Both substituents possess approximately trigonal planar geometry at the CAB atom, and a cone half-angle of 60° was therefore used. Probe positions were generated along 72 uniformly spaced azimuthal directions (5° increments) around the cone. The azimuthal reference direction (0°) was defined by the projection of the CAB to CMB vector onto the plane perpendicular to the cone axis, with increasing angle defined in the direction of the projected CAB to CMA vector. Along each azimuthal direction, probe atoms were placed at 0.05 Å intervals from 0.05 to 2.00 Å from CAB, resulting in 40 sampling positions per ray and 2,880 probe positions per tetrapyrrole.

Signal values from locally filtered cryo-EM maps were sampled at each probe position using ChimeraX (*28*). The resulting values were converted into angle-distance matrices, where the radial coordinate corresponded to distance from CAB and the angular coordinate corresponded to azimuthal position around the cone.

One-dimensional angular density profiles were also extracted at distances corresponding to the expected terminal-atom positions of formyl (approximately 1.2 Å from CAB) and vinyl (approximately 1.35 Å from CAB) substituents. Signal values were averaged across narrow radial windows spanning 1.15-1.25 Å and 1.30-1.40 Å to reduce noise and improve visualization of angular trends.

<u>Förster coupling calculations</u>

Excitation energy transfer rates between antenna chlorophylls were estimated using a Förster-type point dipole approximation. Chlorophylls were represented as transition dipoles defined by vectors between the NB and ND atoms of each tetrapyrrole ring, and inter-pigment distances were approximated using the coordinates of the Mg atoms.

Pairwise coupling strengths were calculated according to:

$$k_{ij} \propto \frac{C_{dd}\,\kappa_{ij}^{2}}{n^{4}R_{ij}^{6}}$$

where $R_{ij}$ is the distance between pigments $i$ and $j$, $\kappa_{ij}^2$ is the orientation factor between transition dipoles, and $n$ is the refractive index (set to 1.55). The orientation factor $\kappa^2$ was computed from normalized dipole vectors and the unit vector connecting pigment centers. Distances were converted from Å to nm prior to calculation.

All pigments were treated as chlorophyll *d*, and a single prefactor ($C_{dd} = 30$) was used to approximate the combined effects of transition dipole strength and spectral overlap. This value was estimated based on reported values for chlorophyll *a* and the similar photophysical properties of chlorophyll *d*. Because all pigments in the calculation were of the same type, this prefactor acts as a uniform scaling factor and does not affect relative coupling strengths.

To ensure applicability of the point dipole approximation, calculations were restricted to antenna chlorophylls. Chlorophylls associated with the electron transfer chain were excluded due to their close spatial proximity and strong electronic coupling, where the dipole approximation is known to break down.

For each structure, full pairwise coupling matrices were computed between antenna chlorophylls. To focus on the most relevant excitation energy transfer pathways, pigment pairs were filtered prior to visualization based on coupling magnitude. Specifically, only interactions with calculated coupling strengths ≥0.04 (arbitrary scaled units) were retained, allowing suppression of weak couplings that are more sensitive to minor geometric variations and are less likely to contribute substantially to excitation energy transfer.

Visualization of coupling networks was performed in PyMOL by representing chlorophyll centers as nodes and pairwise couplings as lines connecting Mg atoms. Line thickness was scaled according to coupling magnitude, such that stronger interactions are represented by thicker connections.

Phylogenetic analyses

A genome-wide species phylogeny for *A. marina* strains (NCBI BioProject PRJNA649288) and *A. thomasi* RCC1774 (NCBI accession NZ_PQWO00000000) was reconstructed with IQ-TREE version 2.0 for a concatenation of 1283 protein sequences from single-copy orthologs identified by OrthoFinder v2.2.7. A maximum likelihood tree was constructed according to the JTT+F+R10 model of sequence evolution selected by the Bayesian information criterion (BIC) in ModelFinder, outgroup-rooted with *Cyanothece* sp. PCC 7425 (NCBI: GCA_000022045.1), and ultrafast bootstrap replicated with 1000 replicates. A similar approach was used for reconstructing PsaB trees for aligned amino acid positions 250-530, 250-475 and 481-530, respectively. A neighbor net network was inferred for an alignment of *A. marina* PsaX alleles with SplitsTree version 4.14.4 (www.splitstree.org).

### Supplementary Text

**Text S1.** Time-resolved fluorescence and global analysis of datasets

The fluorescence emission data on MBIC PSI in the literature are inconsistent. Early fluorescence studies performed at 80 K revealed a dominant emissive band at 730 nm (*35*). However, the spectral shape of its emission profile was strongly dependent on the excitation wavelength. Those studies showed that upon excitation of the chlorophyll Soret band two additional peaks appear at 670 and 700 nm with amplitudes comparable to the main 730-nm band. Subsequent fluorescence excitation studies have shown that those two "blue" bands are likely associated with fluorescence from free chlorophyll *a* and *d* impurities due to disruption during PSI purification or subsequent sample handling. The spectroscopically cleanest emission spectrum with the most pronounced 730 nm band was obtained upon excitation at 490 nm. This wavelength preferentially excites α-carotene. Any carotenoid with comparable conjugation does not have a detectable fluorescence from the lowest excited state. Thus, the chlorophyll *d* emission that was recorded resulted from carotenoid-to-chlorophyll *d* excitation energy transfer, a process that completely bypasses free chlorophylls (*35*). Subsequent attempts at recording fluorescence emission at 77 K with excitation of the Soret band of chlorophyll *d* demonstrated progress in better sample handling as evidenced by much smaller 670 and 700 nm bands (*36*). Most recently, 77 K fluorescence spectra of PSI from MBIC show a clean 730 nm band without any evidence of blue bands, though the spectra were obtained after direct excitation of α-carotene at 490 nm (*37*). Taken together, these studies show that even a minuscule amount of free Chls in the sample can drastically alter fluorescence emission spectra, especially if wavelengths directly exciting chlorophyll Soret bands are utilized. It also indicates that the fluorescence yield of intact PSI from this species has a very low value, much smaller than those of free chlorophylls *a* or *d*.

Since the time-resolved fluorescence (TRF) setup used in this work utilizes 410 nm for excitation, it is expected that the contribution of free chlorophyll emission could be observed to some degree in all studied PSI samples. However, considering that recent TRF studies of PSI from MBIC demonstrated that fluorescence decay from intact PSI has very short-lived dynamics compared to fluorescence decay of free chlorophyll *d*/*a*, those contaminating signals could be easily separated in the spectro-kinetic analysis of the overall TRF signal (*37*). Consequently, kinetic analyses of TRF datasets would allow for more precise reconstruction of expected steady-state fluorescence spectra of those PSI complexes, free of signals associated with unbound chlorophyll *d*/*a*.

A summary of the TRF studies of PSI from MBIC, MU03, MU05 and FH6 strains is shown in **fig. S6**. TRF was recorded upon excitation at 410 nm at 77 K and the data were globally analyzed with kinetic models depicted in panels presenting SAS – species associated spectra (left column) which represent spectral profiles resulting from the fitting procedure and decaying with associated lifetimes (see legends). If the anticipated fitting model of the TRF map is correctly predicted, those profiles should represent fluorescence spectra associated with specific molecular species (i.e., different spectral forms of chlorophyll *d*). Global analysis of the TRF data for MBIC PSI (**fig. S6A**) revealed two spectro-kinetic components that are simultaneously populated with 410 nm excitation and decay independently with indicated lifetimes. The 6.5-ns decay lifetime is associated with a fluorescence maximum at 704 nm and is characteristic of free chlorophyll *d* (*38*). This fraction is not involved in excitation energy transfer within PSI and is likely due to impurities of free pigments. The faster SAS component, revealed as a sharp band with a fluorescence maximum at 732 nm, has a decay lifetime of 50.7 ps and closely resembles the fluorescence spectrum of bulk chlorophyll *d* molecules in the intact MBIC PSI reported recently (*37*). The quality of this fit in the time delay domain is shown in **fig. S6B**, which shows the raw fluorescence decay trace recorded at 732 nm and accompanying fitted curve resulting from global analysis. Considering the global analysis results, the expected spectrum of fluorescence emission of the MBIC PSI could be estimated solely as the 50.7 ps SAS. In **fig. S6C** this spectrum is overlaid with steady-state absorption, indicating that the 732-nm emission is associated with a pool of chlorophylls absorbing at 728 nm. The global analysis of TRF for MU03 PSI is shown in **fig. S6D** and **E**. A simple but convincing fitting model consists of simultaneous excitation and independent decay of two spectral components which are both associated with chlorophylls bound to PSI. One is faster, with a lifetime comparable to MBIC (50.7 ps, **fig. S6A**), and the other is slower, with a lifetime of 371 ps. Whereas the 50.7-ps fluorescence decay component in MBIC PSI was associated with a single fluorescence maximum at 732 nm, the 57-ps fluorescence decay component of MU03 PSI exhibits two maxima – one at 730 nm (similar to MBIC PSI) and a second of similar intensity at 746 nm. These two bands indicate that the network of chlorophyll *d* molecules localized near $P_{740}$ in the electron transfer chain could comprise two spectrally distinct pools of chlorophyll *d* molecules, only one of which is also present in MBIC PSI and the other of which is unique to MU03 PSI. The slower fluorescence decay component in MU03 PSI is broad, with a maximum at 760 nm, and is evidently associated with fluorescence from the lowest energy chlorophyll *d* molecules. This implies that the chlorophyll *d* molecules that give rise to the 371-ps fluorescence decay component are more weakly involved in the process of excitation trapping compared to the 57-ps component.

The global analysis results of TRF for MU05 PSI is presented in **fig. S6G** and **H**. The analysis utilized the same fitting model as for MU03 though a third component was needed to account for a small fraction of free chlorophyll *d* molecules (**fig. S6G**). The two SAS associated with intact PSI have decay lifetimes of 44.9 ps and 395 ps. The faster component appears as a sharp band, probably analogous to the ~745-nm fluorescence maximum in the 50.7 ps component of MU03 PSI. The longer-lived 395-ps component is hypsochromically shifted compared to its counterpart in MU03 (760 nm, 371 ps) and its maximum appears at 755 nm. Comparison of the steady-state absorption spectrum and expected fluorescence emission (with contributions from bulk and red chlorophylls) for MU05 PSI is shown in **fig. S6I.**

Global analysis of TRF for FH6 PSI (**fig. S6J** and **K**) shows two simultaneously and independently decaying components with lifetimes of 29 ps and 390 ps that have corresponding fluorescence maxima at 747 and 765 nm, respectively. Thus, these components are slightly lower in energy and faster than the corresponding components in MU05 PSI. Reconstruction of the expected steady-state fluorescence spectrum of FH6 PSI indicates that a majority is due to emission from red chlorophyll *d* molecules. Moreover, comparison to other PSI complexes with emissive red states clearly indicate that the largest contribution of fluorescence from this molecular species in the overall PSI emission is indeed unique to FH6 PSI.

Together, the TRF data show that PSI from all the strains exhibit a fast fluorescence decay component ~30-50 ps. This component can conceptually be split into contributions from two fluorescence maxima: ~730 nm and 745 nm. MBIC PSI exhibits only the ~730 nm contribution, whereas MU03 PSI exhibits contributions from both ~730 and ~745 nm. MU05 and FH6 PSI both lack the ~730 nm contribution and instead exhibit only the ~745 nm contribution. MU03, MU05, and FH6 all additionally contain a slow component (~400 ps lifetime) at ~760 nm.

**Text S2.** Cryo-EM models and assignments of chlorophylls and carotenoids

*Subunit composition.* The MU03 and MU05 PSI structures each contain 11 subunits. In MU03, these are PsaA, PsaB, PsaC, PsaD, PsaE, PsaF, PsaI, PsaJ, PsaK, and PsaL. In MU05, these are PsaA, PsaB, PsaC, PsaD, PsaE, PsaF, PsaI, PsaK, PsaL, and PsaX. Thus, MU05 PSI lacks PsaJ. Because the *psaJ* gene is present in MU05, we suspect that this subunit was lost during protein isolation. Consistent with this interpretation, the peripheral subunits PsaF and PsaX also exhibit reduced occupancy in MU05 relative to the remainder of the complex. Cofactors coordinated by PSI subunits are listed in **table S6**.

*Tetrapyrrole analyses and assignments.* Pigment analysis by HPLC demonstrated that nearly all chlorophyll molecules in PSI from the different *A. marina* strains are chlorophyll *d*, with chlorophyll *a* comprising only ~1–3% of the total tetrapyrroles (**fig. S9**). All antenna chlorophylls were therefore modeled as chlorophyll *d*.

Due to the consistent observation of ~1% chlorophyll *a*, previous spectroscopic studies proposed that it could be present within the electron transfer chain of *A. marina* PSI (*35*, *39*, *40*). The high resolution of the MU03 and MU05 PSI structures provided an opportunity to directly evaluate this possibility. The PSI electron transfer chain contains six tetrapyrrole cofactors corresponding to the special pair $P_{740}$ ($P_A$ and $P_B$), the "accessory" chlorophylls $A_{-1A}$ and $A_{-1B}$, and the primary electron acceptors $A_{0A}$ and $A_{0B}$ (*11*, *41*). In *A. marina* PSI, $P_A$ contains chlorophyll *d′* (*35*) and the $A_0$ sites contain pheophytin *a* molecules (*9*), leaving $P_B$, $A_{-1A}$, and $A_{-1B}$ as the only sites that could plausibly contain either chlorophyll *a* or chlorophyll *d*. Structurally, chlorophyll *d* differs from chlorophyll *a* by replacement of the C3 vinyl substituent with a formyl group (**fig. S10A**). Because the cryo-EM maps in the electron transfer chain region reach local resolutions of ~1.6 Å, we investigated whether these substituents could be distinguished directly from the cryo-EM density.

To assess tetrapyrrole identity, we adapted the cone-scan method described previously (*33*, *34*). Probe atoms were distributed in a conical sampling volume extending from the C3 substituent position (**fig. S10A**), and cryo-EM density values were sampled throughout the expected spatial region occupied by either a formyl or vinyl group (**fig. S10B-M**). As controls, the analysis included $P_A$, which can be confidently assigned as chlorophyll *d′* based on pigment analysis (**fig. S9**) and the characteristic prime stereochemistry of the $13^2$ methoxycarbonyl substituent, and the two $A_0$ sites, which can be confidently assigned as pheophytin *a* because pheophytin *d* has not been detected in *A. marina* and because the cryo-EM maps clearly lack density corresponding to a central Mg atom.

The resulting density distributions did not reveal a clear feature that distinguished the C3 formyl moiety of the known chlorophyll *d′* in site $P_A$ from the C3 vinyl moieties of the known pheophytin *a* molecules in the $A_0$ sites (**fig. S10**), suggesting that these two groups cannot be distinguished in the cryo-EM data using this approach. Thus, the $P_B$, $A_{-1A}$, and $A_{-1B}$ sites were all modeled as the bulk chlorophyll type, chlorophyll *d*. Interestingly, however, the density profiles of all six C3 moieties were highly reproducible between the MU03 and MU05 PSI (**fig. S10**) despite their independent determination and differences elsewhere in the structures. Although the absolute density values are not directly comparable because the maps are independently scaled, the overall shapes, relative intensities, and directional features of the profiles are remarkably similar between strains. Furthermore, the $A_{-1B}$ site exhibited consistently stronger density in the C3 substituent region than the other five tetrapyrroles, including the known vinyl-containing pheophytin *a* molecules, in both MU03 and MU05 PSI (**fig. S10**). The origin of this feature remains unclear. Several sites also exhibit preferred directions of density extending from the C3 substituent region, often accompanied by weaker density in the opposite direction. Given the high local resolution of these maps, these features likely reflect the geometry of the substituent and neighboring atoms, including the hydrogen atoms attached to $C3_1$, rather than alternate conformations.

One possible explanation for the limited discriminatory power of the analysis is that the exceptionally high local resolution may actually hinder differentiation of formyl- and vinyl-substituents. Theoretically, differences arising from negative or partial negative charge become less pronounced as cryo-EM maps approach higher resolution (*42*). Consequently, the electrostatic-potential signatures of C=O and C=C groups may be more similar than expected in maps approaching atomic resolution, limiting direct structural discrimination even when the substituents themselves are well resolved. Additional studies combining spectroscopy and targeted mutagenesis could help to definitively determine whether chlorophyll *a* can occupy electron transfer chain sites in *A. marina* PSI.

*Carotenoid analysis.* Pigment analysis showed that nearly all carotenoids in *A. marina* PSI from each strain are α-carotene, with only minor populations of zeaxanthin and an unidentified carotenoid detected by HPLC (**fig. S11A-C**). Because carotenoids participate in light harvesting, photoprotection, and structural stabilization in photosystems, we evaluated whether multiple carotenoid species could be identified structurally. Most carotenoid sites exhibited one well-resolved ring and a second substantially more flexible ring, limiting confident assignment of carotenoid identity at many positions. Nevertheless, one carotenoid site in the MU03 PSI structure, corresponding to site J15, displayed additional density consistent with a hydroxylated carotenoid ring (**fig. S11D**), such as that present in zeaxanthin (**fig. S11C**). To further evaluate this assignment, we quantitatively compared map signal extending from the carotenoid ring at multiple positions and found that the density more closely resembled vectors corresponding to known substituents than those terminating in hydrogen atoms. Accordingly, we tentatively modeled this site as zeaxanthin, although it could also correspond to the unidentified carotenoid observed by HPLC. Notably, the unidentified carotenoid eluted earlier than α-carotene during HPLC separation, suggesting that it is less hydrophobic and may therefore contain a polar substituent such as a hydroxyl group.

The reduced map quality at this site in MU05 is likely related to the absence of the nearby PsaJ subunit, which appears to destabilize several peripheral regions of the complex. In MU03, the tentatively assigned zeaxanthin is positioned adjacent to a chlorophyll *d* molecule such that the carotenoid hydroxyl group lies within hydrogen-bonding distance (~2.8 Å) of the oxygen atom of the chlorophyll C3 formyl group. Although the same chlorophyll environment is generally preserved in MU05, the corresponding region additionally contains the sidechain of PsaB-Leu427, which may sterically hinder this interaction. In MU03, this position is instead occupied by the smaller residue PsaB-Ser429, whose hydroxyl group is oriented away from the carotenoid-binding pocket, potentially permitting accommodation of the carotenoid hydroxyl group. Additional studies will be required to determine whether this carotenoid influences excitation-energy transfer, photoprotection, or the spectral properties of nearby chlorophyll *d* molecules.

**Text S3.** Map and model details near the region that may correspond to the fluorescence component at ~760 nm

The peripheral region of PsaB implicated in formation of the ~760 nm fluorescence component (*5*) exhibits substantially lower local resolution than the remainder of the PSI structures in MBIC (*9*), MU03, and MU05, indicating pronounced structural flexibility in this region. In the previously determined MBIC PSI structure, large portions of this region could not be modeled, including segments centered near PsaB residues ~300 and ~480, which lacked 30 and 46 amino acids, respectively. In the MU03 PSI structure, the same regions remain partially unresolved but improved local map quality allowed substantially more residues to be modeled, reducing the missing segments to 6 and 19 amino acids, respectively. Nevertheless, many residues in these regions remain poorly resolved and their precise conformations should be interpreted cautiously. MU03 and MU05 exhibit highly similar sequences in these regions, and comparison of MBIC, MU03, MU05, and NIES suggests that the PsaX-associated region is generally more ordered in PsaX-containing PSI complexes, building upon and strengthening previous similar hypotheses (*5*, *9*).

To improve interpretation of this region, we performed local focused refinements of this region for both MU03 and MU05 PSI. Structural interpretation was additionally guided by comparison to PSI from *Thermosynechococcus vestitus*, which is more analogous to MU05, and *Synechocystis* sp. PCC 6803 (hereafter *Synechocystis*), which is more analogous to MU03. In the region centered near PsaB residue 480, the MU03 map clearly supports the presence of only a single chlorophyll corresponding to site B31, whereas *Synechocystis* PSI contains two chlorophylls, B31 and B32. No additional density corresponding to a second chlorophyll was observed in MU03, although loss of a weakly occupied chlorophyll cannot be completely excluded.

Interpretation of the region centered near PsaB residue 300 was aided by stronger conservation of chlorophyll-coordinating residues between MU03 and *Synechocystis*. Chlorophylls B18 and B19 are coordinated by conserved histidine residues in both systems, and these residues are also conserved in MBIC. Assignment of chlorophyll B40 is less certain because its associated looping region remains largely unresolved in MU03 PSI and entirely unresolved in MBIC PSI. In *Synechocystis* PSI, B40 is coordinated indirectly through interactions involving a phosphatidylglycerol molecule and the adjacent PsaB loop. In MU03 PSI, weak density consistent with the phosphate group of a phosphatidylglycerol molecule is present and was tentatively modeled. Despite uncertainty in the associated protein loop, the chlorophyll ring corresponding to B40 is clearly resolved (**fig. S12D**). Interestingly, the B40 tetrapyrrole ring in MU03 is substantially more parallel to B18 and B19 than in *Synechocystis* PSI, resulting in a more trimer-like chlorophyll arrangement in MU03 PSI. Thus, it seems like a good candidate for the 755-nm TRF component.

Several nearby chlorophylls in MU03 PSI, including B9, B17, and B18, also exhibit relatively weak density, although their axial ligands are conserved with *Synechocystis* and their assignments are considered reliable. Outside this region, the MU03 PSI structure is generally well resolved except for a short flexible loop in PsaK centered near residue 52, a feature commonly observed in cyanobacterial PSI structures. PsaF and PsaJ, which often exhibit reduced occupancy in cyanobacterial PSI structures, are comparatively well resolved in the MU03 PSI structure.

The MU05 PSI structure is generally better resolved in the PsaX-associated region than MU03. However, as described in **text S2**, PsaJ is entirely absent and both PsaF and PsaX exhibit reduced occupancy, although the remaining regions could still be modeled reliably. As in MU03, a short loop in PsaK centered near residue ~52 could not be modeled.

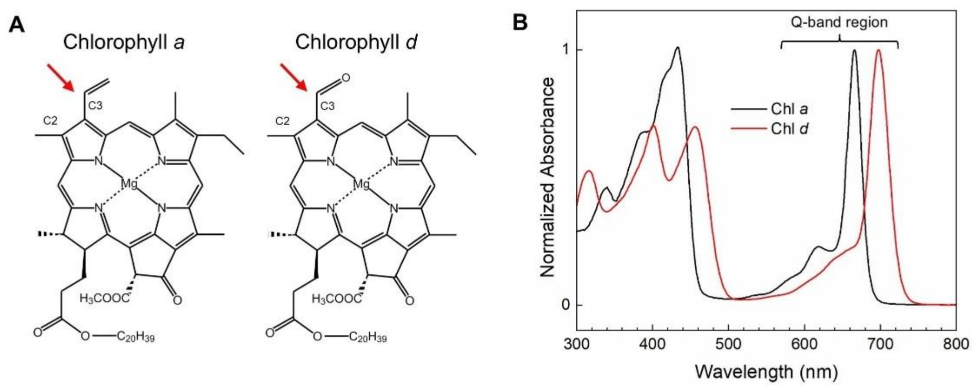

**Fig. S1.** Structures and absorption spectra of chlorophyll *a* and chlorophyll *d*. (**A**) Structures of chlorophylls *a* and *d*. Red arrows point to the single substituent difference between the two chlorophyll types. (**B**) Absorption spectra of chlorophyll *a* and *d* in methanol. Spectra are normalized to the $Q_y$ maximum.

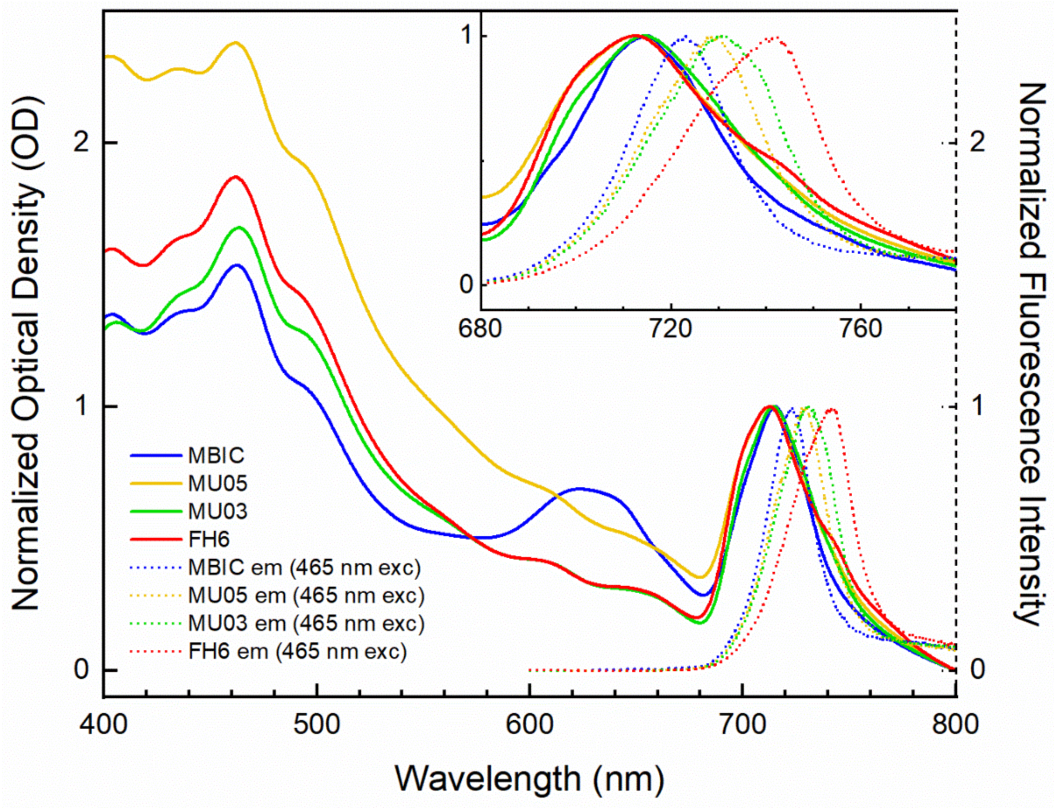

**Fig. S2.** Whole cell absorption and fluorescence emission spectra of *A. marina* strains.

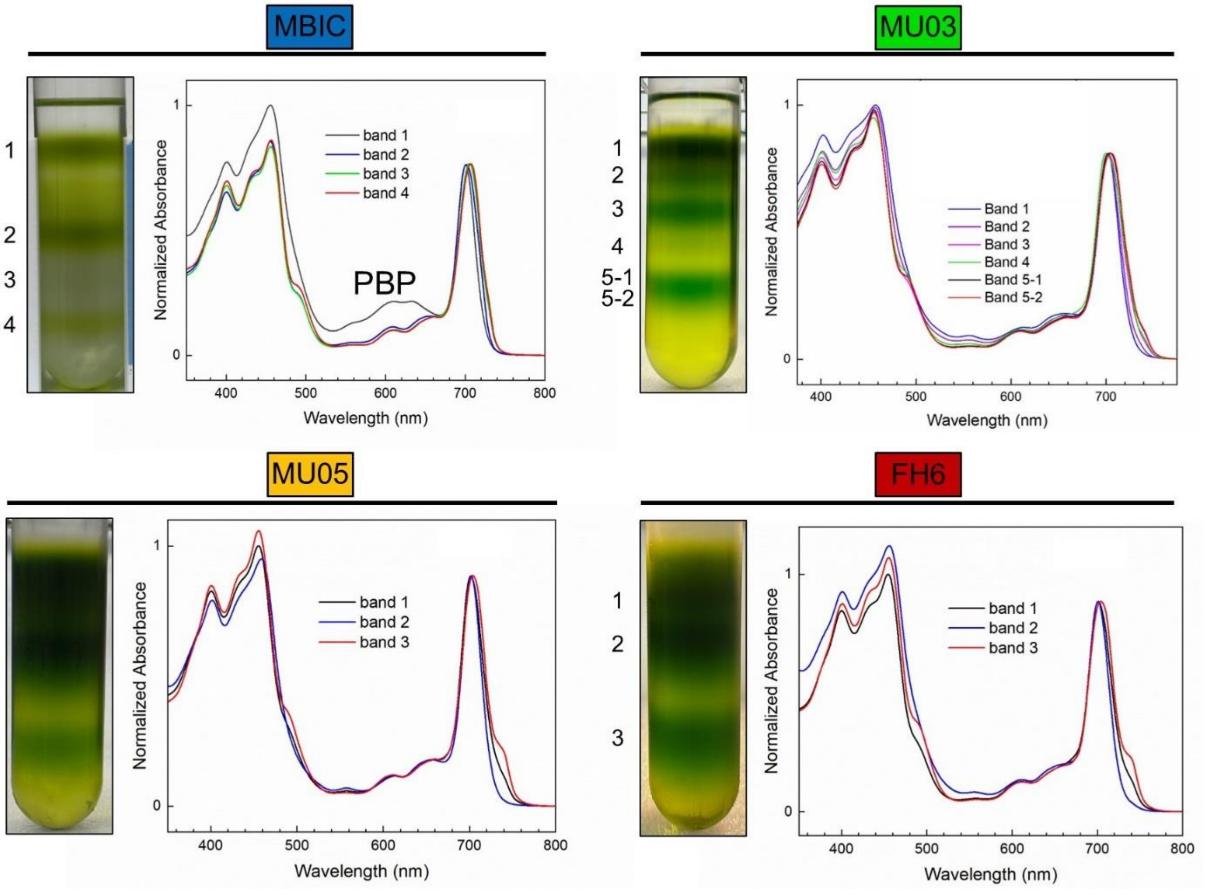

**Fig. S3.** Separation of photosynthetic complexes from *A. marina* cells. For each strain, the trehalose gradient, and the corresponding absorption spectra from each band are shown. MBIC is the only of these strains containing phycobiliproteins. This contribution to the absorption (band 1) is labeled “PBP”.

**PsbA (D1)**

```
MBIC PsbA1 MTTVLQ---RRESASAWERFCSFITSTNNRLYIGWFGVLMIPTLLTAVTCFVIAFIGAPP 57
MU05 PsbA1 MTTVLQ---RRESASAWERFCSFITSTNNRLYIGWFGVLMIPTLLTAVTCFVIAFIGAPP 57
FH6 PsbA1  MTTVLQ---RRESASAWERFCSFITSTNNRLYIGWFGVLMIPTLLTAVTCFVIAFIGAPP 57
MU03 PsbA  MTTVLQ---RRESASAWERFCSFITSTNNRLYIGWFGVLMIPTLLTAVTCFVIAFIGAPP 57
MU05 PsbA3 MSTTFQTRTRFSLFSIWDQFCDWITSTQNRLYIGWFGVLMIPTLLVSAITFILAWVAAPS 60
FH6 PsbA3  MSTTFQTRTRFSLFSIWDQFCDWITSTQNRLYIGWFGVLMIPTLLVSAITFILAWVAAPS 60
MBIC PsbA2 MSTTFQTPSRLPTVSAWDQFCEWITSTHNRLYVGWFGLLMIPSLFVSAITFMLAWVAAPS 60
MU05 PsbA2 MSTTLQARSRLPIYSIWEQFCEWITSTQHRLYIGWFGVVMIPTLFVAAITFLLAWVAAPS 60
FH6 PsbA2  MSTTLQARSRLPIYSIWEQFCEWITSTQHRLYIGWFGVVMIPTLFVAAITFLLAWVAAPS 60
           *:*.:*   *    * *::**.:****::***:****::***:*:.:.  *::*::.**

MBIC PsbA1 VDIDGIREPVAGSLLYGNNIITGAVVPSSNAIGLHLYPIWEAASLDEWLYNGGPYQLIIF 117
MU05 PsbA1 VDIDGIREPVAGSLLYGNNIITGAVVPSSNAIGLHLYPIWEAASLDEWLYNGGPYQLIIF 117
FH6 PsbA1  VDIDGIREPVAGSLLYGNNIITGAVVPSSNAIGLHLYPIWEAASLDEWLYNGGPYQLIIF 117
MU03 PsbA  VDIDGIREPVAGSLLYGNNIITGAVVPSSNAIGLHLYPIWEAASLDEWLYNGGPYQLVIF 117
MU05 PsbA3 VDMDGIREPVIGSLMSGNNVITAAVIPSSAAIGLHLYPLWEATSLDEWLYNGGPYQLIIL 120
FH6 PsbA3  VDMDGIREPVIGSLMSGNNVITAAVIPSSAAIGLHLYPLWEATSLDEWLYNGGPYQLIIL 120
MBIC PsbA2 VDMEGIREPIISSLLGGSNVITAAVIPTSAAIGLHLYPLWEATSMDEWLYNGGPYQLIIL 120
MU05 PsbA2 VDMDGIREPLIGSLMGGNNVITAAVIPTSAAIGLHLYPLWEATSVDEWLYNGGPYQMIIL 120
FH6 PsbA2  VDMDGIREPLIGSLMGGNNVITAAVIPTSAAIGLHLYPLWEATSVDEWLYNGGPYQMIIL 120
           **::*****: .**: *.*:**.**:*:* ********:***:*:***********::*:

MBIC PsbA1 HYMIGCICYLGRQWEYSYRLGMRPWICVAYSAPLAATYSVFLIYPLGQGSFSDGMPLGIS 177
MU05 PsbA1 HYMIGCICYLGRQWEYSYRLGMRPWICVAYSAPLAATYSVFLIYPLGQGSFSDGMPLGIS 177
FH6 PsbA1  HYMIGCICYLGRQWEYSYRLGMRPWICVAYSAPLAATYSVFLIYPLGQGSFSDGMPLGIS 177
MU03 PsbA  HYMIGCICYLGRQWEYSYRLGMRPWICVAYSAPLAATYSVFLIYPLGQGSFSDGMPLGIS 177
MU05 PsbA3 HFLIAIWAYLGRQWELSYRIGMRPWIAMAFSAPVAAATAVLLVYLMGQGSFSESLPLGIA 180
FH6 PsbA3  HFLIAIWAYLGRQWELSYRIGMRPWIAMAFSAPVAAATAVLLVYLMGQGSFSESLPLGIA 180
MBIC PsbA2 HFLIAIWTYLGRQWELSYRLGMRPWIAMAFSAPVAAATAVLLVYPMGQGSFSEGLPLGIS 180
MU05 PsbA2 HFLIAIWAYLGRQWEFSYRIGMRPWIAMAFSAPVAAATAVLLVYPMGQGSFSEGLPLGIA 180
FH6 PsbA2  HFLIAIWAYLGRQWEFSYRIGMRPWIAMAFSAPVAAATAVLLVYPMGQGSFSEGLPLGIA 180
           *::*.   ******* ***:******.:*:***:**: :*:*:* :******:.:****:

MBIC PsbA1 GTFNFMFVFQAEHNILMHPFHMFGVAGVLGGSLFAAMHGSLVSSTLVRETTEGESANYGY 237
MU05 PsbA1 GTFNFMFVFQAEHNILMHPFHMFGVAGVLGGSLFAAMHGSLVSSTLVRETTEGESANYGY 237
FH6 PsbA1  GTFNFMFVFQAEHNILMHPFHMFGVAGVLGGSLFAAMHGSLVSSTLVRETTEGESANYGY 237
MU03 PsbA  GTFNFMFVFQAEHNILMHPFHMMGVAGVLGGSLFAAMHGSLVSSTLVRETTEGESANYGY 237
MU05 PsbA3 GTFHFMMAVQADHNILMHPFHMLGVVGVFGGAFLSAMHGSLVASSLVQETSSLESVNTGY 240
FH6 PsbA3  GTFHFMMAVQADHNILMHPFHMLGVVGVFGGAFLSAMHGSLVASSLVQETSSLESVNTGY 240
MBIC PsbA2 GTFHFMMAVQAEHNILMHPFHMLGVVGVFGGAFLSAMHGSLVTSSLVQETSSLKSVNTGY 240
MU05 PsbA2 GTFHFMMAVQADHNILMHPFHMLGVVGVFGGAFLSAMHGSLVASSLVEETSSLESVNTGY 240
FH6 PsbA2  GTFHFMMAVQADHNILMHPFHMLGVVGVFGGAFLSAMHGSLVASSLVEETSSLESVNTGY 240
           ***:**:..**:**********:**.**:**::::*******:*:**.**:. :*.* **

MBIC PsbA1 KFGQEEETYNIVAAH-GYFGRLIFQYASFSNSRSLHFFLGAWPVVCIWLTAMGISTMAFN 296
MU05 PsbA1 KFGQEEETYNIVAAH-GYFGRLIFQYASFSNSRSLHFFLGAWPVVCIWLTAMGISTMAFN 296
FH6 PsbA1  KFGQEEETYNIVAAH-GYFGRLIFQYASFSNSRSLHFFLGAWPVVCIWLTAMGISTMAFN 296
MU03 PsbA  KFGQEEETYNIVAAH-GYFGRLIFQYASFSNSRSLHFFLGAWPVVCIWLTAMGISTMAFN 296
MU05 PsbA3 KFGQQGLWPRQRRNWAISL--KAMPVIVLSS----------------------------- 269
FH6 PsbA3  KFGQQGLWPRQRRNWAISL--KAMPVIVLSS----------------------------- 269
MBIC PsbA2 KFGQQEATYNLLAGHAGYLGRLFIPDIAFRNSRSIHFLLAVLPTIGIWFAALGIGTMAFN 300
MU05 PsbA2 KFGQQKATYNLLAGHAGYLGRLFIPVIAFRNSRSIHFLLATLPTIGIWFAALGIGTIAFN 300
FH6 PsbA2  KFGQQKATYNLLAGHAGYLGRLFIPVIAFRNSRSIHFLLATLPTIGIWFAALGIGTIAFN 300
           ****:    .        :    :    : .

MBIC PsbA1 LNGFNFNHSIVDSQGNVVNTWADVLNRANLGFEVMHERNAHNFPLDLAAGESAPVALTAP 356
MU05 PsbA1 LNGFNFNHSIVDSQGNVVNTWADVLNRANLGFEVMHERNAHNFPLDLAAGESAPVALTAP 356
FH6 PsbA1  LNGFNFNHSIVDSQGNVVNTWADVLNRANLGFEVMHERNAHNFPLDLAAGESAPVALTAP 356
```

```
MU03 PsbA  LNGFNFNHSIVDSQGNVVNTWADVLNRANLGFEVMHERNAHNFPLDLAAGESAPVALTAP 356
MU05 PsbA3 ------------------------------------------------------------ 269
FH6 PsbA3  ------------------------------------------------------------ 269
MBIC PsbA2 LNGFNFNHSLLDSSGRPIRTEADLLNRATMGLQVMHSVNAHHFSLTLASTESKEIPT-IP 359
MU05 PsbA2 LNGFNFNHSLLDSSGRTIRTEADLLNRATMGLQVMHSVNAHNFPLTLASTDSMETPT-IP 359
FH6 PsbA2  LNGFNFNHSLLDSSGRTIRTEADLLNRATMGLQVMHSVNAHNFPLTLASTDSMETPT-IP 359

MBIC PsbA1 VING 360
MU05 PsbA1 VING 360
FH6 PsbA1  VING 360
MU03 PsbA  VING 360
MU05 PsbA3 ---- 269
FH6 PsbA3  ---- 269
MBIC PsbA2 IMTS 363
MU05 PsbA2 IISS 363
FH6 PsbA2  IISS 363
```

**PsbD (D2)**

```
MBIC PsbD1 MTIAVGRAQERGWFDVLDDWLKRDRFVFIGWSGILLFPCAFLSIGGWFTGTTFVTSWYTH 60
MBIC PsbD2 MTVALGRVQERGWFDVLDDWLKRDRFVFIGWSGLLLFPCAFLSIGGWFTGTTFVTSWYTH 60
MU03 PsbD1 --IAVGRAQERGWFDVLDDWLKRDRFVFIGWSGILLFPCAFLSIGGWFTGTTFVTSWYTH 58
MU03 PsbD2 --IAVGRAQERGWFDVLDDWLKRDRFVFIGWSGILLFPCAFLSIGGWFTGTTFVTSWYTH 58
MU05 PsbD  MTIAVGRAQERGWFDVLDDWLKRDRFVFIGWSGILLFPCAFLSVGGWFTGTTFVTSWFTH 60
FH6 PsbD   MTIAVGRAQERGWFDVLDDWLKRDRFVFIGWSGILLFPCAFLSVGGWFTGTTFVTSWFTH 60
             :*:**.*************************:*********:*************:**

MBIC PsbD1 GLASSYLEGANFLTVAVSTPADSLGHSLLLLWGPEAQGDFTRWCQLGGLWNFTTLHGVFG 120
MBIC PsbD2 GLASSYLEGCNFLTAAVSTPADSMGHSLLLLWGPEARGDFTRWCQLGGMWNFVTLHGAFG 120
MU03 PsbD1 GLASSYLEGANFLTVAVSTPADSLGHSLLLLWGPEAQGDFTRWCQLGGLWNFTTLHGVFA 118
MU03 PsbD2 GLASSYLEGANFLTVAVSTPADSLGHSLLLLWGPEAQGDFTRWCQLGGLWNFTTLHGVFA 118
MU05 PsbD  GLASSYLEGANFLTVAVSTPADSLGHSLLLLWGPEAQGDFTRWCQLGGLWNFTTLHGVFG 120
FH6 PsbD   GLASSYLEGANFLTVAVSTPADSLGHSLLLLWGPEAQGDFTRWCQLGGLWNFTTLHGVFG 120
           *********.****.********:************:***********:***.****.*.

MBIC PsbD1 LIGFMLRQFEIARLVGVRPYNAVAFSGPIAVYVSVFLMYPLGQSSWFFAPSWGVTSIFRF 180
MBIC PsbD2 LIGFMLRQFEIARLVNVRPYNAVAFSGPIAVFVSVFLMYPLGQSSWFFAPSWGVASIFRF 180
MU03 PsbD1 LIGFMLRQFEVARLIGVRPYNAVAFSGPIAVFVSVFLMYPLGQSSWFFAPSWGVTSIFRF 178
MU03 PsbD2 LIGFMLRQFEVARLIGVRPYNAVAFSGPIAVFVSVFLMYPLGQSSWFFAPSWGVTSIFRF 178
MU05 PsbD  LIGFMLRQFEVARLIGVRPYNAVAFSGPIAVYVSVFLMYPLGQSSWFFAPSWGVTSIFRF 180
FH6 PsbD   LIGFMLRQFEVARLIGVRPYNAVAFSGPIAVYVSVFLMYPLGQSSWFFAPSWGVTSIFRF 180
           **********:***:.***************:**********************:*****

MBIC PsbD1 LLFAQGFHNLTLNPFHMMGVAGILGGALLCAIHGATVENTLFEDGQDANTFAAFTPTQAE 240
MBIC PsbD2 LLFVQGFHNLTLNPFHMMGVAGILGGALLCAIHGATVENTLFEDTKDANTFSGFSPTQSE 240
MU03 PsbD1 LLFAQGFHNLTLNPFHMMGVAGILGGALLCAIHGATVENTLFEDGQDANTFAAFSPTQSE 238
MU03 PsbD2 LLFAQGFHNLTLNPFHMMGVAGILGGALLCAIHGATVENTLFEDGQDANTFAAFSPTQSE 238
MU05 PsbD  LLFAQGFHNLTLNPFHMMGVAGILGGALLCAIHGATVENTLFEDGQDANTFAAFSPTQAE 240
FH6 PsbD   LLFAQGFHNLTLNPFHMMGVAGILGGALLCAIHGATVENTLFEDGQDANTFAAFSPTQAE 240
           ***.**************************************** :*****:.*:***:*

MBIC PsbD1 ETYSMVTANRFWSQIFGIAFSNKRWLHFFMLFVPVTGLWASAIGLVGIALNMRAYDFVSQ 300
MBIC PsbD2 ETYSMVTANRFWSQIFGIAFSNKRWLHFFMLFVPVTGLWASAIGLVGIALNMRAYDFVSQ 300
MU03 PsbD1 ETYSMVTANRFWSQIFGIAFSNKRWLHFFMLFVPVTGLWASAIGLVGIALNMRAYDFVSQ 298
MU03 PsbD2 ETYSMVTANRFWSQIFGIAFSNKRWLHFFMLFVPVTGLWASAIGLVGIALNMRAYDFVSQ 298
MU05 PsbD  ETYSMVTANRFWSQIFGIAFSNKRWLHFFMLFVPVTGLWASAIGLVGIALNMRAYDFVSQ 300
FH6 PsbD   ETYSMVTANRFWSQIFGIAFSNKRWLHFFMLFVPVTGLWASAIGLVGIALNMRAYDFVSQ 300
           ************************************************************

MBIC PsbD1 EIRAAEDPEFETFYTKNILLNEGLRAWMAPQDQIHENFIFPEEVLPRGNAL 351
MBIC PsbD2 EIRAAEDPEFETFYTKNILLNEGLRAWMAPQDQIHENFVFPEEVLPRGNAL 351
MU03 PsbD1 EIRAAEDPEFETFYTKNILLNEGLRAWMAPQDQIHENFIFP---------- 339
```

```
MU03 PsbD2 EIRAAEDPEFETFYTKNILLNEGLRAWMAPQDQIHENFIFP---------- 339
MU05 PsbD  EIRAAEDPEFETFYTKNILLNEGLRAWMAPQDQIHENFIFPEEVLPRGNAL 351
FH6 PsbD   EIRAAEDPEFETFYTKNILLNEGLRAWMAPQDQIHENFIFPEEVLPRGNAL 351
           **************************************:**
```

### PsbB (CP47)

```
MBIC PsbB MGLPWYRVHTVVLNDPGRLLSVHLMHTALVSGWAGSMALYELAKYDPSDPVLNPMWRQGT 60
MU05 PsbB MGLPWYRVHTVVLNDPGRLLSVHLMHTALVSGWAGSMALYELAKYDPSDPVLNPMWRQGT 60
MU03 PsbB MGLPWYRVHTVVLNDPGRLLSVHLMHTALVSGWAGSMALYELAKYDPSDPVLNPMWRQGT 60
FH6 PsbB  MGLPWYRVHTVVLNDPGRLLSVHLMHTALVSGWAGSMALYELAKYDPSDPVLNPMWRQGT 60
          ************************************************************

MBIC PsbB FVMPVMTRIGVTHSWSGWTVTGEPWVTQPGILGAHLNFFSYEGVILMHILAAGLFFLAAV 120
MU05 PsbB FVMPVMTRIGVTHSWSGWTVTGEPWVTQPGILGAHLNFFSYEGVILMHILAAGLFFLAAV 120
MU03 PsbB FVMPVMTRIGVTHSWSGWTVTGEPWVSQSGILGAHLNFFSYEGVILMHILASGLFFLAAV 120
FH6 PsbB  FVMPVMTRIGVTHSWSGWTVTGEPWVSQPGILGAHLNFFSYEGVILMHILAAGLFFLAAV 120
          **************************:* **********************:********

MBIC PsbB WHWINWDLDIYYPDGSSEPASDWPKIFGLHLLTLGIVCFGFGSLHLTGILGPGMWVSDPY 180
MU03 PsbB WHWINWDLDVYYPEGSAESASDWPKIFGLHLLTLGIICFGFGSLHLTGILGPGMWVSDPY 180
FH6 PsbB  WHWINWDLDIYYPDGSSEPASDWPKIFGLHLLTLGIVCFGFGSLHLTGILGPGMWVSDPY 180
MU05 PsbB WHWINWDLDIYYPDGSSEPASDWPKIFGLHLLTLGIVCFGFGSLHLTGILGPGMWVSDPY 180
          *********:***:**:* ****************:***********************

MBIC PsbB GLTGHVQGVSPDWRPFAFDPYNPTGLVTHHISAGIALIIGGIFHTVSRPSERLYNALSMG 240
MU05 PsbB GLTGHVQGVSPDWRPFAFDPYNPTGLVTHHISAGIALIIGGIFHIVSRPSERLYNALSMG 240
MU03 PsbB GLTGHVQGVSPDWRPLAFDPYNPTGLVTHHISAGIALIIGGIFHTVSRPSERLYSALSMG 240
FH6 PsbB  GLTGHVQGVSPDWRPFAFDPYNPTGLVTHHISAGIALIIGGIFHTVSRPSERLYNALSMG 240
          ***************:**************************** *********.*****

MBIC PsbB NVETVLSSSVAFVAAAAFVMVGTMWYGSATTPIELFGPTRYQWDSGYFQTEIQRRVQSGQ 300
MU05 PsbB NVETVLSSSVAFVAAAAFVMVGTMWYGSATTPIELFGPTRYQWDSGYFQTEIQRRVQSGQ 300
MU03 PsbB NVETVLSSSIAFVAAAAFVMVGTMWYGSATTPIELFGPTRYQWDSGYFQQEIQRRVQSGQ 300
FH6 PsbB  NVETVLSSSVAFVAAAAFVMVGTMWYGSATTPIELFGPTRYQWDSGYFQTEIQRRVQSGQ 300
          *********:*************************************** **********

MBIC PsbB TWDQIPEKLVFYDYIGNSPAKGGLFRTGAMNSGDGIARAWEGHPTFTDSEGRELFVRRMP 360
MU05 PsbB TWDQIPEKLVFYDYIGNSPAKGGLFRTGAMNSGDGLARAWEGHPTFTDSEGRELFVRRMP 360
MU03 PsbB DWTQIPEKLVFYDYIGNSPAKGGLFRTGAMNSGDGLARSWEGHPTFTDSEGRELFVRRMP 360
FH6 PsbB  TWDQIPEKLVFYDYIGNSPAKGGLFRTGAMNSGDGLARSWEGHPTFTDSEGRELFVRRMP 360
           * ********************************:**:*********************

MBIC PsbB NFFETFPVVLTDKDGVVRADIPFRRAESRYSFEQKGVSVSFEGGTLNGQTFTDAPSVKKY 420
MU05 PsbB NFFETFPVILTDKDGVVRADIPFRRAESRYSFEQKGVSVSFEGGTLNGQTFTDAPSVKKY 420
MU03 PsbB NFFETFPVVLTDADGVLRADIPFRRAESRYSFEQKGVTVSFEGGTLNGQTFTDAPSVKKY 420
FH6 PsbB  NFFETFPVILTDKDGVIRADIPFRRAESRYSFEQKGVTVSFEGGTLNGQTFTDAPSVKKY 420
          ********:*** ***:********************:**********************

MBIC PsbB ARKAQLGEPFEFDRETLGSDGVFRTSTRGWFAFSHSCYALLFFFGHWWHGARTIFKDVFE 480
MU05 PsbB ARKAQLGEPFEFDRETLGSDGVFRTSTRGWFAFSHSCYALLFFFGHWWHGARTIFKDVFE 480
MU03 PsbB ARKAQLGEPFEFDTETLNSDGVFRTSTRGWFAFSHSCYALLFFFGHWWHGARTIFKDVFE 480
FH6 PsbB  ARKAQLGEPFEFDRETLGSDGVFRTSTRGWFAFSHSCYALLFFFGHWWHGARTIFKDVFE 480
          ************* ***.******************************************

MBIC PsbB GVDPNLPEEQVEWGVFQKVGDTTTRA 506
MU05 PsbB GVDPNLPEEQVEWGVFQKVGDTSTRA 506
MU03 PsbB GVDPNLPEEQVEWGVFQKVGDTSTRA 506
FH6 PsbB  GVDPNLPEEQVEWGVFQKVGDTSTRA 506
          **********************:***
```

### PsbC (CP43)

```
MBIC PsbC MKVCALGWHPKTKSMKTSSSLRRFYHVETPFNPSAAGYDRATTGYGWWAGNARLTDLSGQ 60
```

```
MU05 PsbC --------------------------METPFNPSAAGYDRATTGYGWWAGNARLTDLSGQ 34
MU03 PsbC --------------------------METPFNPSAAGYDRASTGYGWWAGNARLTDLSGQ 34
FH6 PsbC  --------------------------METPFNPSAAGYDRATTGYGWWAGNARLTNLSGQ 34
                                    :**************:*************:****

MBIC PsbC LTGAHIAHAGMITFWAGAMTLFEVSHFIPEKPMYEQGSILLAHLAAEGFGVGPGGEVIST 120
MU05 PsbC LTGAHIAHCGMITFWAGAMTLFEVSHFIPEKPMYEQGSILLAHLAAEGFGVGAGGEVIST 94
MU03 PsbC LTGAHIAHCGMITFWAGAMTLFEVSHFIPEKPMYEQGSILLAHLAAEGFGVGAGGEVIST 94
FH6 PsbC  LTGAHIAHCGMITFWAGAMTLFEVSHFIPEKPMYEQGSILLAHLAAEGFGVGAGGEVIST 94
          ********.******************************************* *******

MBIC PsbC YPYFVIGAIHLIASAVLGFGGLYHTFRGPAKFEDYSDWWGYDWEDKEKMMQILGIHLIFL 180
MU05 PsbC YPYFVIGAIHLIASAVLGFGGLYHTFRSPAKFEDFSDYWGYDWEDKEKMMQILGIHLIFL 154
MU03 PsbC YPYFVIGAIHLIASAVLGFGGLYHTFRGPAKFEDYSDWWGYDWEDKEKMMQILGIHLIFL 154
FH6 PsbC  YPYFVIGALHLIASAVLGFGGLYHTFRSPAKFEDFSDYWGYDWEDKEKMMQILGIHLIFL 154
          ********:******************.******:**:**********************

MBIC PsbC GIGALAFAAKAMFFGGLYDPWAPGGGNVRLITNPTWNLGTFLGYITRSPWGEGGWIVSVN 240
MU05 PsbC GIGALAFAAKAMFFGGLYDPWAPGGGNVRLITNPTWNLGTFLGYITRSPWGEGGWIVSVN 214
MU03 PsbC GIGALAFAAKAMFFGGLYDPWAPGGGNVRLITNPTWNLGTFLGYITRSPWGEGGWIVSVN 214
FH6 PsbC  GIGALLFAAKAMFFGGLYDPWAPGGGNVRLITNPTWNLGTFLGYITRSPWGEGGWIVSVN 214
          ***** ******************************************************

MBIC PsbC NLEDVVGGHLLVGVHYIFGGVFHILVKPWGWVRRAYVWSGEAYLSYSLGALYMCGMIAVG 300
MU05 PsbC NLEDVVGGHLLVGVHYIFGGVFHILVKPWGWVRRAYVWSGEAYLSYSLGALYMCGMIAVG 274
MU03 PsbC NLEDVVGGHLLVGVHYIFGGVFHILVKPWGWVRRAYVWSGEAYLSYSIGALYMCGMIAVG 274
FH6 PsbC  NLEDVVGGHLLVGVHYIFGGVFHILVKPWGWVRRAYVWSGEAYLSYSIGALYMCGMIAVG 274
          ***********************************************:************

MBIC PsbC YVWFNNTVYPSEFYGPTAAEASQAQAMTFLIRDQRLGANIASAQGPTGLGKYLMRSPSGE 360
MU05 PsbC YVWFNNTVYPSEFYGPTAAEASQAQAMTFLIRDQRLGANIASAQGPTGLGKYLMRSPSGE 334
MU03 PsbC YIWFNNTVYPSEFYGPTAAEASQAQAMTFLIRDQRLGANIASAQGPTGLGKYLMRSPSGE 334
FH6 PsbC  YIWFNNTVYPSEFYGPTAAEASQAQAMTFLIRDQRLGANIASAQGPTGLGKYLMRSPSGE 334
          *:**********************************************************

MBIC PsbC IIFGGETMRFWDFRGPWLEPLRGPNGLDLNKLRNDIQPWQARRAAEYMTHAPLGALNSVG 420
MU05 PsbC IIFGGETMRFWDFRGPWLEPLRGPNGLDLNKLRNDIQPWQARRAAEYMTHAPLGALNSVG 394
MU03 PsbC IIFGGETMRFWDFRGPWLEPLRGPNGLDLNKLRNDIQPWQARRAAEYMTHAPLGALNSVG 394
FH6 PsbC  IIFGGETMRFWDFRGPWLEPLRGPNGLDLNKLRNDIQPWQARRAAEYMTHAPLGALNSVG 394
          ************************************************************

MBIC PsbC GVATEINSVNYVSPRSWLSTSHFCLAFFFFVGHIWHSGRARAAAAGFEKGIERKTEYALS 480
MU05 PsbC GVATEINSVNYVSPRSWLSTSHFCLAFFFFVGHIWHSGRARAAAAGFEKGIERKTEYALS 454
MU03 PsbC GVATEINSVNYVSPRSWLSTSHFCLAFFFFVGHIWHSGRARAAAAGFEKGIERKTEYALS 454
FH6 PsbC  GVATEINSVNYVSPRSWLSTSHFCLAFFFFVGHIWHSGRARAAAAGFEKGIERKTEYALS 454
          ************************************************************

MBIC PsbC LPDIDATSVD 490
MU05 PsbC LPDIDATAVD 464
MU03 PsbC LPDIDATSVD 464
FH6 PsbC  LPDIDATAVD 464
          *******:**
```

**Fig. S4.** Multiple sequence alignments of PSII subunits comparing *A. marina* strains. This alignment excludes one incomplete PsbA sequence from MU05 and one incomplete PsbD sequence from FH6. Sequences were aligned using Clustal Omega (*43*).

### PsaA

```
MBIC PsaA MTTSPGGPETKGRTAEVDINPVSASLEVAGKPGHFNKSLSKGPQTTTWIWNLHALAHDFD 60
MU03 PsaA MTISPGGPETKGRTAEVDINPVPASLEVAGKPGHFNKTLSRGPQTTTWIWNLHALAHDFD 60
MU05 PsaA MTTSPGGPETKGRTAEVDINPVKASLEVAGKPGHFNKSLSKGPQTTTWIWNLHALAHDFD 60
FH6 PsaA  MTNSPGGPETKGRTAEVDINPVKASLEVAGKPGHFNKSLSKGPQTTTWIWNLHALAHDFD 60
          ** ******************* **************:**:*******************

MBIC PsaA TQTNDLEEISRKIFSAHFGHLSIIFVWISGMIFHAARFSNYYAWLADPLGNKPSAHVVWP 120
MU03 PsaA TQTNDLEEISRKIFSAHFGHLSIIFVWISGMIFHAARFSNYYAWLADPFGNKPSAHVVWP 120
MU05 PsaA TQTNDLEEISRKIFSAHFGHLAIVFVWISGMIFHAARFSNYYAWLADPLGNKPSAHVVWP 120
FH6 PsaA  TQTNDLEEISRKIFSAHFGHLAIVFVWISGMIFHAARFSNYYAWLADPLGNKPSAHVVWP 120
          *********************:*:************************:***********

MBIC PsaA IVGQDILNADVGNGFRGVQITSGLFHILRGAGMTDPGELYSAAIGALVAAVVMMYAGYYH 180
MU03 PsaA IVGQDILNADVGNGFHGIQITSGLFHILRGAGMTAPIELYSTSIGALVAAAVTMYAGYFH 180
MU05 PsaA IVGQDILNADVGNGFHGIQITSGLFHILRGAGMTAPIELYSTSIGALVAAAVTMYAGYFH 180
FH6 PsaA  IVGQDILNADVGNGFHGIQITSGLFHILRGAGMTAPIELYSTSIGALVAAAVTMYAGYFH 180
          ***************:*:**************** * ****::*******.* *****:*

MBIC PsaA YHKKAPKLEWFQNAESTMTHHLIVLLGLGNLAWTGHLIHVSLPVNKLLDSGVAPQDIPIP 240
MU03 PsaA YHKKAPKLEWFQNAEATIGGHLIVLLGLGNLAWTGHLIHVSLPINKMLDSGISPEDIPIP 240
MU05 PsaA YHKKAPKLEWFQNAEATIGGHLINLLGLGNLAWTGHLIHVSLPINKMLDSGMAPEDIPIP 240
FH6 PsaA  YHKKAPKLEWFQNAEATIGGHLINLLGLGNLAWTGHLIHVALPINKMLDSGMAPEDIPIP 240
          ***************:*:  *** ****************:**:**:****::*:*****

MBIC PsaA HEFLFDNGFMADLYPSFAQGLMPYFTLNWGAYSDFLTFKGGLDPTTGGLWMTDIAHHHLA 300
MU03 PsaA HEYLLDKAFMADLYPSFAQGLAPYFTLNWGVYSDFLTFKGGLDPQTGGLWMTDIAHHHLA 300
MU05 PsaA HEYLLDKAFMADLYPSFAQGLAPYFTLNWGVYSDFLTFKGGLDPQTGGLWMTDIAHHHLA 300
FH6 PsaA  HEYLLDKAFMADLYPSFAQGLTPFFTLNWGVYSDFLTFKGGLDPQTGGLWMTDIAHHHLA 300
          **:*:*:.************* *:******.************* ***************

MBIC PsaA LAVMYIIAGHMYRTNWGIGHSMKEIMESHKGPFTGEGHKGLYEVLTTSWHAQLAINLATW 360
MU03 PsaA LAVMYIIAGHMYRTNWGIGHHMKEIMEAHKGPFTGEGHKGLYEVLTTSWHAQLAINLSCW 360
MU05 PsaA LAVMYIIAGHMYRTNWGIGHHMREIMDAHKGPFTGEGHKGLYDVLTTSWHAQLAINLACW 360
FH6 PsaA  LAVMYIIAGHMYRTNWGIGHHMREIMDAHKGPFTGEGHKGLYDVLTTSWHAQLAINLACW 360
          ******************** *:***::**************:**************: *

MBIC PsaA GSFSIIVAHHMYAMPPYPYLATDYGTQLNLFVHHMWIGGFLIVGGAAHAAIFMVRDYDPA 420
MU03 PsaA GSFSIIVAHHMYAMPPYPYLATDYATQLNLFVHHMWLGGFYIVGGAAHAAIFMVRDYDPE 420
MU05 PsaA GSFSIVCAHHMYAMPPYPYLATDYATQLNLFVHHMWLGGFYIVGGAAHAAIFMVRDYDPA 420
FH6 PsaA  GSFSIVCAHHMYAMPPYPYLATDYATQLNLFVHHMWLGGFYIVGGAAHAAIFMVRDYDPA 420
          *****: *****************.***********:*** ****************** 

MBIC PsaA VNQNNVLDRMLRHRDTIISHLNWVCIFLGFHSFGLYIHNDNMRSLGRPQDMFSDTAIQLQ 480
MU03 PsaA MNQNNVLDRMLRHRDTIISHLNWVCIFLGFHSFGLYIHNDEMRSLGRPQDMFSDTAIQLQ 480
MU05 PsaA MNQNNVLDRMLRHRDTIIAHLNWVCLFLGFHSFDLYIHNDEMRSLGRPQDMFSDTAIQLQ 480
FH6 PsaA  MNQNNVLDRMLRHRDTIIAHLNWVCLFLGFHSFDLYIHNDEMRSLGRPQDMFSDTAIQLQ 480
          :*****************:******:*******.******:*******************

MBIC PsaA PIFSQWVQNLQANVAGTIRAPLAEGASSLAWGGDPLFVGGKVAMQHVSLGTADFMIHHIH 540
MU03 PsaA PIFAQWVQNLQANVAGTIRAPLAEGATSLAWGGDPLIVSGKVAMQHVSLGTADFMIHHIH 540
MU05 PsaA PIFAQWVQNLQANVAGTIRAPLAEGATSLAWGGDPMFVGGKVAMQHVSLGTADFMIHHIH 540
FH6 PsaA  PIFAQWVQNLQANVAGTIRAPLAEGATSLAWGGDPMFVGGKVAMQHVSLGTADFMIHHIH 540
          ***:**********************:********::*.*********************

MBIC PsaA AFQIHVTVLILIKGVLYARSSRLIPDKANLGFRFPCDGPGRGGTCQSSGWDHIFLGLFWM 600
MU03 PsaA AFQIHVTILILMKGVLYARSSRLIPDKASLGFRFPCDGPGRGGTCQSSGWDHVFLGLFWM 600
MU05 PsaA AFQIHVTVLILLKGVLYARSSRLIPDKANLGFRFPCDGPGRGGTCQSSGWDHVFLGLFWM 600
FH6 PsaA  AFQIHVTVLILLKGVLYARSSRLIPDKASLGFRFPCDGPGRGGTCQSSGWDHVFLGLFWM 600
          *******:***:****************.***********************:*******

MBIC PsaA YNCISIVNFHFFWKMQSDVWGAANANGGVNYLTAGNWAQSSITINGWLRDFLWAQSVQVI 660
```

```
MU03 PsaA YNCISIVNFHFFWKMQSDVWGSANADGAINYITAGNWAQSSITINGWLRDFLWAQSVQVI 660
MU05 PsaA YNCISIVNFHFFWKMQSDVWGTANASGGINYLTAGNWAQSSITINGWLRDFLWAQSVQVI 660
FH6 PsaA  YNCISIVNFHFFWKMQSDVWGTANASGGINYLTAGNWAQSSITINGWLRDFLWAQSVQVI 660
          *********************:***.*.:**:****************************

MBIC PsaA NSYGSALSAYGILFLGAHFIWAFSLMFLFSGRGYWQELIESIVWAHSKLKIAPAIQPRAM 720
MU03 PsaA NSYGSALSAYGILFLGAHFIWAFSLMFLFSGRGYWQELIESIVWAHGKLKIAPAIQPRAM 720
MU05 PsaA NSYGSALSAYGILFLGAHFIWAFSLMFLFSGRGYWQELIESIVWAHSKLKIAPAIQPRAM 720
FH6 PsaA  NSYGSALSAYGILFLGAHFIWAFSLMFLFSGRGYWQELIESIVWAHSKLKIAPAIQPRAM 720
          **********************************************.*************

MBIC PsaA SITQGRAVGLGHYLLGGIVTSWSFYLARILALG 753
MU03 PsaA SITQGRAVGLGHYLLGGIVTSWSFYLARILALG 753
MU05 PsaA SITQGRAVGLGHYLLGGIVTSWSFYLARILALG 753
FH6 PsaA  SITQGRAVGLGHYLLGGIVTSWSFYLARILALG 753
          *********************************
```

**PsaB**

```
MBIC PsaB MATKFPSFSQDLAQDPTTRRIWYGIATVHDFETHDGMTEENLYQKIFATHFGHLSIIFLW 60
MU03 PsaB MATKFPSFSQDLAQDPTTRRIWYGIATTHDFETHDGMTEENLYQKIFATHFGHLSIIFLW 60
MU05 PsaB MATKFPSFSQDLAQDPTTRRIWYGIATVHDFETHDGMTEENLYQKIFATHFGHLSIIFLW 60
FH6 PsaB  MATKFPSFSQDLAQDPTTRRIWYGIATVHDFETHDGMTEENLYQKIFATHFGHLSIIFLW 60
          ***************************.********************************

MBIC PsaB SAGHLFHVAWQGNFEQWIQDPLTIRPIAHAIWDPHLGDAATQAFTQAGASGPVDLCYSGL 120
MU03 PsaB AAGLLFHVAWQGNFEQWIQDPLTVRPIAHAIWDPHLGQSATEAFTQAGASGPVDICYSGV 120
MU05 PsaB TAGLLFHVAWQGNFEQWIQDPLTVRPIAHAIWDPHLGDAATQAFTQAGASGPVDLCYSGV 120
FH6 PsaB  TAGLLFHVAWQGNFEQWIQDPLTVRPIAHAIWDPHLGDAATQAFTQAGASGPVDLCYSGV 120
          :** *******************:*************::**:************:****:

MBIC PsaB YQWWYTIGMRTNGDLYIGSVFLMIVAAVMLFAGWLHLQPKFRPSLAWFRDAESQMNHHLA 180
MU03 PsaB YHWWYTIGMRTNADLYSGSIFLMVVASVMLFAGWLHLQPKFRPSLAWFRDAESQMNHHLA 180
MU05 PsaB YQWWYTIGMRTNGDLYSGSIFLMVVAAIMLFAGWLHLQPKFRPSLAWFRDAESQMNHHLA 180
FH6 PsaB  YHWWYTIGMRTNGDLYSGSIFLMVVAAVMLFAGWLHLQPKFRPSLAWFRDAESQMNHHLA 180
          *:**********.*** **:***:**::********************************

MBIC PsaB VLFGASSLGWTGHLIHVAIPEARGQHVGWDNFLSTMPHPAGLAPFFTGRWGVYAQNPDTA 240
MU03 PsaB VLFGASSLGWTGHLIHVAIPEARGQHVGWDNFLSTMPHPAGLAPFFTGRWAVYAQNPDTA 240
MU05 PsaB VLFGASSLGWTGHLIHVAIPEARGQHVGWDNFLSTMPHPAGLAPFFTGRWAVYAQNPDTA 240
FH6 PsaB  VLFGASSLGWTGHLIHVAIPEARGQHVGWDNFLSTMPHPAGLAPFFTGRWAVYAQNPDTT 240
          **************************************************.********:

MBIC PsaB GHIFGTSEGAGTAIITFIGGFHPQTEALWLTDIAHHHLAIAVMYIIAGHMYRTQFGIGHS 300
MU03 PsaB GHIFGSSEGSGTAILTFIGGFHPQSEALWLTDIAHHHLAIAVMYIIAGHMYRTQFGIGHS 300
MU05 PsaB GHIFGTSEGAGTAILTFMGGFHPQTEALWLTDIAHHHLAIAVLYIVAGHMYRTQFGIGHS 300
FH6 PsaB  GHLFGTSEGAGTAILTFVGGFHPQTEALWLTDIAHHHLAIAVLYIVAGHMYRTQFGIGHS 300
          **:**:***:****:**:******:*****************:**:**************

MBIC PsaB MKEILEAHTPPSGMLGDAHKGLYDTYNESLHFQLGFHLAALGVITSVVAQHMYSLPSYAF 360
MU03 PsaB MKEILEAHTPPSGILGDAHKGLYDTYNESLHFQLGFHLAALGVITSVVAQHMYSLPSYAF 360
MU05 PsaB MKEIMDAHRDP--WYGATLQGLYDTYNESLHFQLAFHLAALGVITEVVAQHMYSLPSYAF 358
FH6 PsaB  MKEIMDAHRDP--WYGATLEGLYDTYNESLHFQLAFHLAALGVITEVVAQHMYSLPSYAF 358
          ****::**  *    * : :**************.**********.**************

MBIC PsaB ISQDHVTQAALYTHHQYIAGILAIGAFAHGGIFFVRDYDPERNKNNVLARALEHKEAIIS 420
MU03 PsaB IAEDYATQAALYTHHQYIAGILAIGAFAHGGIFFVRDYDPERNKNNVLARALAHKEAIIS 420
MU05 PsaB ISQDYVTQAALYTHHQYIGGFLALGAYAHGGIFFVRDYDPERNKGNALSRLLEHKEAIIS 418
FH6 PsaB  ISQDYVTQAALYTHHQYIGGFLALGAYAHGGIFFVRDYDPERNKGNALSRLLEHKEAIIS 418
          *::*:.************.*:**:**:*****************.*.*:* * *******
```

```
MBIC PsaB HLSWVSMFSGFHTLGVYVHNDTVVAFGTPEKQILVEPIFAQWIQAAHGKLLLGFETLLSN 480
MU03 PsaB HLSWVSMFSGFHTLGVYVHNDVVVAFGTPEKQILVEPIFAQWIQAAHGKLLLGFETLLSN 480
MU05 PsaB HLSWVSMFLGFHTLDLYVHNDVVVAFGTPEKQILPEPIFAEWVQAAHGKLLLGLDSLLSN 478
FH6 PsaB  HLSWVSMFLGFHTLDLYVHNDVVVAFGTPEKQILPEPIFAEWVQAAHGKLLLGLDSLLSN 478
          ******** *****.:*****.************ *****:*:**********:::****

MBIC PsaB PNGLAYNPPNISPDVFVPGWVEAMNNPVIGPFMSQGPGDFLVHHGIAFSLHVTVLICVKG 540
MU03 PsaB PNGLASNPPNISPDVFVPGWVEAMNNPVLGPFMSQGPGDFLVHHGIAFSLHVTVLICVKG 540
MU05 PsaB PQSIASTAWPNYGDVWLPGWLDAVNGA-NTPFLNIGPGDFLVHHGIAFSIHVTVLICVKG 537
FH6 PsaB  PQSIAATAWPNYGDVWLPGWLDAVNGS-STPFLNIGPGDFLVHHGIAFSIHVTVLICVKG 537
          *:.:* .      **::***::*:*.    **:. **************:**********

MBIC PsaB CLDARGSKLMPDKKDFGYSFPCDGPGRGGTCDISAWDSFYLAFFWMLNTIGWIVFYFNWK 600
MU03 PsaB CLDARGSKLMPDKKDFGYSFPCDGPGRGGTCDISAWDSFYLAFFWMLNTIAWITFYFNWK 600
MU05 PsaB CLDARGSKLMPDKKDFGYSFPCDGPGRGGTCDISAWDSFYLATFWMLNTIGWVTFYFNWK 597
FH6 PsaB  CLDARGSKLMPDKKDFGYSFPCDGPGRGGTCDISAWDSFYLATFWMLNTIGWVTFYFNWK 597
          ****************************************** *******.*:.******

MBIC PsaB HLAIWSGNEAQFNTNSTYLMGWLRDYLWGYSAQLINGYTPFGVNSLSVWAWIFLLGHLCW 660
MU03 PsaB HLAIWAGNEAQFNTNSTYLMGWLRDYLWGYSAQLINGYTPFGVNSLSVWAWIFLLGHFVW 660
MU05 PsaB QLSVWSGNLAQFNNDSTYLMGWLRDYLWGYSAQLINGYTPFGVNSLSVWAWIFLLGHLCW 657
FH6 PsaB  QLSVWSGNLAQFNNDSTYLMGWLRDYLWGYSAQLINGYTPFGVNSLSVWAWIFLLGHLCW 657
          :*::*:** ****.:******************************************: *

MBIC PsaB ATGFLFLISWRGYWQELIETLVWAHQRTPLANLVTWKDKPVALSIVQGRLVGLVHFAVGY 720
MU03 PsaB AIGFLFLISWRGFWQELIETLVWAHQRTPLANLVQWKDKPVALSIVQGRLVGLVHFALGY 720
MU05 PsaB ATGFLFLISWRGYWQELIETLVWAHQRTPLANLVMWKDKPVALSIVQGRLVGLVHFAVGY 717
FH6 PsaB  ATGFLFLISWRGYWQELIETLVWAHQRTPLANLVMWKDKPVALSIVQGRLVGLVHFAVGY 717
          * **********:********************* **********************:**

MBIC PsaB YVTYAAFVIGATAPLG 736
MU03 PsaB VVTYGAFVIGATAPLS 736
MU05 PsaB FVTYGAFVIGATAPLG 733
FH6 PsaB  FVTYGAFVIGATAPLG 733
           ***.**********.
```

**PsaC**

```
MBIC PsaC MSHTVKIYDTCIGCTQCVRACPTDVLEMVPWDGCKAGQIASSPRTEDCVGCKRCETACPT 60
MU03 PsaC MSHTVKIYDTCIGCTQCVRACPLDVLEMVPWDGCKAGQIASSPRTEDCVGCKRCETACPT 60
MU05 PsaC MSHTVKIYDTCIGCTQCVRACPTDVLEMVPWDGCKAGQIASSPRTEDCVGCKRCETACPT 60
FH6 PsaC  MSHTVKIYDTCIGCTQCVRACPTDVLEMVPWDGCKAGQIASSPRTEDCVGCKRCETACPT 60
          ********************** *************************************

MBIC PsaC DFLSIRVYLGAETTRSMGLAY 81
MU03 PsaC DFLSIRVYLGAETSRSMGLAY 81
MU05 PsaC DFLSIRVYLGAETTRSMGLAY 81
FH6 PsaC  DFLSIRVYLGAETSRSMGLAY 81
          *************:*******
```

**PsaD**

```
MBIC PsaD MAETLTGQTPLFGGSTGGLLSSAETEEKYAITWTSPKQQVFEMPTGGAAVMNEGENLLYL 60
MU03 PsaD MADTLTGQTPLFGGSTGGLLSSADTEEKYAITWTSPKQQVFEMPTGGAAVMNEGENLLYL 60
MU05 PsaD MAETLTGQTPLFGGSTGGLLSSAETEEKYAITWTSPKQQVFEMPTGGAAVMNEGENLLYL 60
FH6 PsaD  MAETLTGQTPLFGGSTGGLLSSAETEEKYAITWTSPKQQVFEMPTGGAAVMNEGENLLYL 60
          **:********************:************************************

MBIC PsaD ARKEQCLALGLRQLRTKKIMDYKIYRVLPDGSNTLLHPKDGVFPEKSNEGRAAVNSVARS 120
MU03 PsaD ARKEQCLALGVRQLRTKKIMDYKIYRVLPDGSNTMIHPKDGVFPEKVNEGRQAVGSVARS 120
MU05 PsaD ARKEQCLALGLRQLRTKKIMDYKIYRVLPDGSNTLLHPKDGVFPEKSNEGRTAVNSVARS 120
FH6 PsaD  ARKEQCLALGLRQLRIKKIMDYKIYRVLPDGSNTLLHPKDGVFPEKSNEGRAAVNSVARS 120
```

```
          **********:**** *****************::********** **** **.*****

MBIC PsaD IGENPNPGAIKYTGKKAYD 139
MU03 PsaD IGENPNPGAMKYSGKNPYD 139
MU05 PsaD IGENPNPGAIKYTGKKAYD 139
FH6 PsaD  IGENPNPGAIKYTGKQAYD 139
          *********:**:**: **
```

**PsaE**

```
MBIC PsaE MVQRGSKVRILRPESYWFREIGTVASVDQSGIKYPVIVRFDTCNYNGISGTAAGINTNNF 60
MU03 PsaE MVQRGSKVRILRPESYWFREIGTVASVDQSGIKYPVIVRFETCNYNGISGTAAGVNTNNF 60
MU05 PsaE MVQRGSKVRILRPESYWFREIGTVASVDQSGIKYPVIVRFDTCNYNGISGTAAGVNTNNF 60
FH6 PsaE  MVQRGSKVRILRPESYWFREIGTVASVDKSGIKYPVIVRFDTCNYNGISGTAAGINTNNF 60
          ****************************:***********:*************:*****

MBIC PsaE GVHELEEVEAPKGGKKPAAKP---AAAPKAEG----- 89
MU03 PsaE GLHEVEEVEAPKGGKKKAPAAAKAAAAPKEEAKPADG 97
MU05 PsaE GLHELEEVEAPKAK---AKKP---AA-PKAEG----- 85
FH6 PsaE  GVHELEEVEAPKAK---AKKP---AAAPKAEG----- 86
          *:**:*******.    *      ** ** *.
```

**PsaF**

```
MBIC PsaF MRRLFAVLLVMTLFLGVVPPASADIGGLVPCSESPKFQERAAKARNTTADPNSGQKRFEM 60
MU03 PsaF MRRLFAVLLVMTLWLGAASPALAEIGGLVPCSESPKFQDRATKARNTTADPNSGQKRFEM 60
MU05 PsaF MRRLFAVLLVMTLWLGVVPPASADIGGLVPCSESPKFQERAAKARNTTGDPNSGQKRFEM 60
FH6 PsaF  MRRLFAVLVVLTLWLGVVPPASADIGGLVPCSESPQFQERAAKARNTTGDPESGRKRFEM 60
          ********:*:**:**.. ** *:***********:**:**:******.**:**:*****

MBIC PsaF YSSALCGPEDGLPRIIAGGPMRRAGDFLIPGLFFIYIAGGIGNSSRNYQIANRKKNAKNP 120
MU03 PsaF YSSALCGADDGLPRIIAGGPMSRAGDFLIPGLFFIYIAGGIGNASRNYQIANRKKNPKNP 120
MU05 PsaF YSSALCGPEDGLPRIIAGGPWSRAGDFLIPGLLFIYIAGGIGNASRNYQIANRKKNPKNP 120
FH6 PsaF  YSSALCGADDGLPRIIAGGPWSRAGDFLIPGLLFIYIAGGIGNASRNYQIANRKKNPKNP 120
          ******* :***********  **********:**********:************ ***

MBIC PsaF AMGEIIIDVPLAVSSTIAGMAWPLTAFRELTSGELTVPDSDVTVSPR 167
MU03 PsaF AMGEIIIDVPLALSSTIAALAWPLTAFRELTSGQLTVPDSDVTTSPR 167
MU05 PsaF AMGEIIIDVPLALSSTIAALAWPLKALGELTSGKLTVPDSDVTVSPR 167
FH6 PsaF  AMGEIIIDVPLALSSTIAALAWPLKALGEVTSGKLTVPDSDVTVSPR 167
          ************:*****.:****.*: *:***:*********.***
```

**PsaI**

```
MBIC PsaI MISDILPAIMTPLVVLIGGGAAMTAFFYYVEREG 34
MU03 PsaI MISDILPAIMTPLVCLVGAGTAMTAFFYYVEREG 34
MU05 PsaI MISDILPAIMTPLVVLIGGGAAMTAFFYYVEREG 34
FH6 PsaI  MISDILPAIMTPLVVLIGGGAAMTAFFYYVEREG 34
          ************** *:*.*:*************
```

**PsaJ**

```
MBIC PsaJ MKFFSTAPVIALVFFTLTAGFLVELNRFFPDILFFPY 37
MU03 PsaJ MKFFSTAPVIAFIFFTLTAGFMVELNRFFPDILFFPF 37
MU05 PsaJ MKFFSTAPVIAFIFFTLTAGFLVELNRFFPDILFFPY 37
FH6 PsaJ  MKFFSTAPVIAFVFFTLTAGFLVELNRFFPDILFFPY 37
          ***********::********:**************:
```

**PsaK**

```
MBIC PsaK1 ---------------------MIAANILAIAIAKYSIQYPNVGPSMPAS--NLFGGFSFN   37
MU03 PsaK1 ---------------------MIAANIMGIAIAKYSIQYPNVGPSMPAS--NLFGGFSLT   37
```

```
MU05 PsaK1 ---------------------MIAANILAIAIAKYSIQYPNVGPSMPAS--SLFGGFSFN   37
FH6 PsaK1  ---------------------MIAANIMAIAIAKYSIQYPNVGPSMPAS--SLFGGFSFN   37
MBIC PsaK2 MHPILLTTFESHPWTVNTGILMLFINLLMVFLGRYAIKYPGQGPALPIGVPDSMKDFGVP   60
MU03 PsaK2 MHPILLVAFESHSWTINTGILMLVINLLMVAIGRYTIKYPGQGPALPFNVPDSMKDFGVP   60
MU05 PsaK2 MHPILLTTFESHPWTVNTGILMLFINLLMVFLGRYTIKYAGQGPALPFAVPEFMKDFGIP   60
FH6 PsaK2  MHPILLTTFESHPWTFNTGILMLFINLLMVFLGRYAIKYPGQGPALPIGVPDSMKDFGVP   60
                                *:  *:: : :.:*:*:* . **::*    . : .*..

MBIC PsaK1 AVLGTQVFGHVLGAGAILGLTYLGVL   63
MU03 PsaK1 SVLGTQVFGHILGAGAVLGLTYLGVI   63
MU05 PsaK1 AVLGTQVFGHVLGAGAILGLTYLGVL   63
FH6 PsaK1  AVLGTQVFGHVLGAGAILGLTYLGVL   63
MBIC PsaK2 EMLATGVFAHWIGAGMILGLRSAGAL   86
MU03 PsaK2 EMLATGAFAHWIGAGMILGLRTSGAL   86
MU05 PsaK2 EMLATGVFAHWIGAGMILGLRSAGAL   86
FH6 PsaK2  EMLATGVFAHWIGAGMILGLRSAGAL   86
            :*.* .*.* :*** :***   *.:
```

#### PsaL

```
MBIC PsaL MTMDLVKHGGDPLVGDLATPVNSSGIVKAWINNLPAYRKGMSANARGLEIGMAHGYYLYG 60
MU03 PsaL MTMEMVKHGGDPFVGDLATPVNSSGIVKAWINNLPAYRKGMSANARGLEIGMAHGYYLYG 60
MU05 PsaL MTMDLVTHGGDPFVGDLATPVNSSGIVKAWINNLPAYRKGMSANARGLEIGMAHGYYLYG 60
FH6 PsaL  MTMEMVKHGGDPFVGDLATPVNSSGIVKAWINNLPAYRKGMSANARGLEIGMAHGYYLYG 60
          ***::*.*****:***********************************************

MBIC PsaL PFATSGPIRGTTMSLVSGVLSASAVVIVLSVAIQIYSSVSSAKPLPSVTTADPGSDFGTK 120
MU03 PsaL PFATSGPIRGTQMALVSGVLSASCVIIVLSVAIQLYTSVSVPKPRPSVTTADPGDDFGTK 120
MU05 PsaL PFATSGPIRGTTMSLVSGVLSASAVIIVLSVAIQIYSSVSSAKPHPSVTTADPGSDFATK 120
FH6 PsaL  PFATSGPVRGTTMALVSGVLSASCVIIVLTVAMQLYSSLSVPKPRPSVTTADPGSDFGTK 120
          *******:*** *:*********.*:***:**:*:*:*:*  ** *********.**.**

MBIC PsaL EGWSAVGSGFLIGGCGGAVIAGVLSYAIAAFVG 153
MU03 PsaL EGWSAMGSGFLIGGCGGAVIAGVLSYAIAIFAG 153
MU05 PsaL EGWSAIGSGFLIGGCGGAVIAGVLSYAIAIFAG 153
FH6 PsaL  EGWSAIGSGFLIGGCGGAVIAGVLSYAIAIFAG 153
          *****:*********************** *.*
```

#### PsaM

```
MBIC PsaM MELSDLQIVIALVVALLPALLALNLGSALSK 31
MU03 PsaM MELSDLQIVIALVVALLPALLAVNLGSALSK 31
MU05 PsaM MELSDLQIVIALVVALLPALLALNLGSALSK 31
FH6 PsaM  MELSDLQIVIALVVALLPALLALNLGTALSK 31
          **********************:***:****
```

#### PsaX

```
MU05_PsaX MDNTGKSPWLTLLLIWSGLVVAFLFLSNLQIG 32
FH6_PsaX  MNKTTKNPWPTLPLIWSGIGILAAIWITLQIG 32
          *::* *.** ** *****: :   :  .****
```

**Fig. S5.** Multiple sequence alignments of PSI subunits comparing *A. marina* strains. Note that MBIC and MU03 do not contain genes for *psaX*. Sequences were aligned using Clustal Omega (*43*).

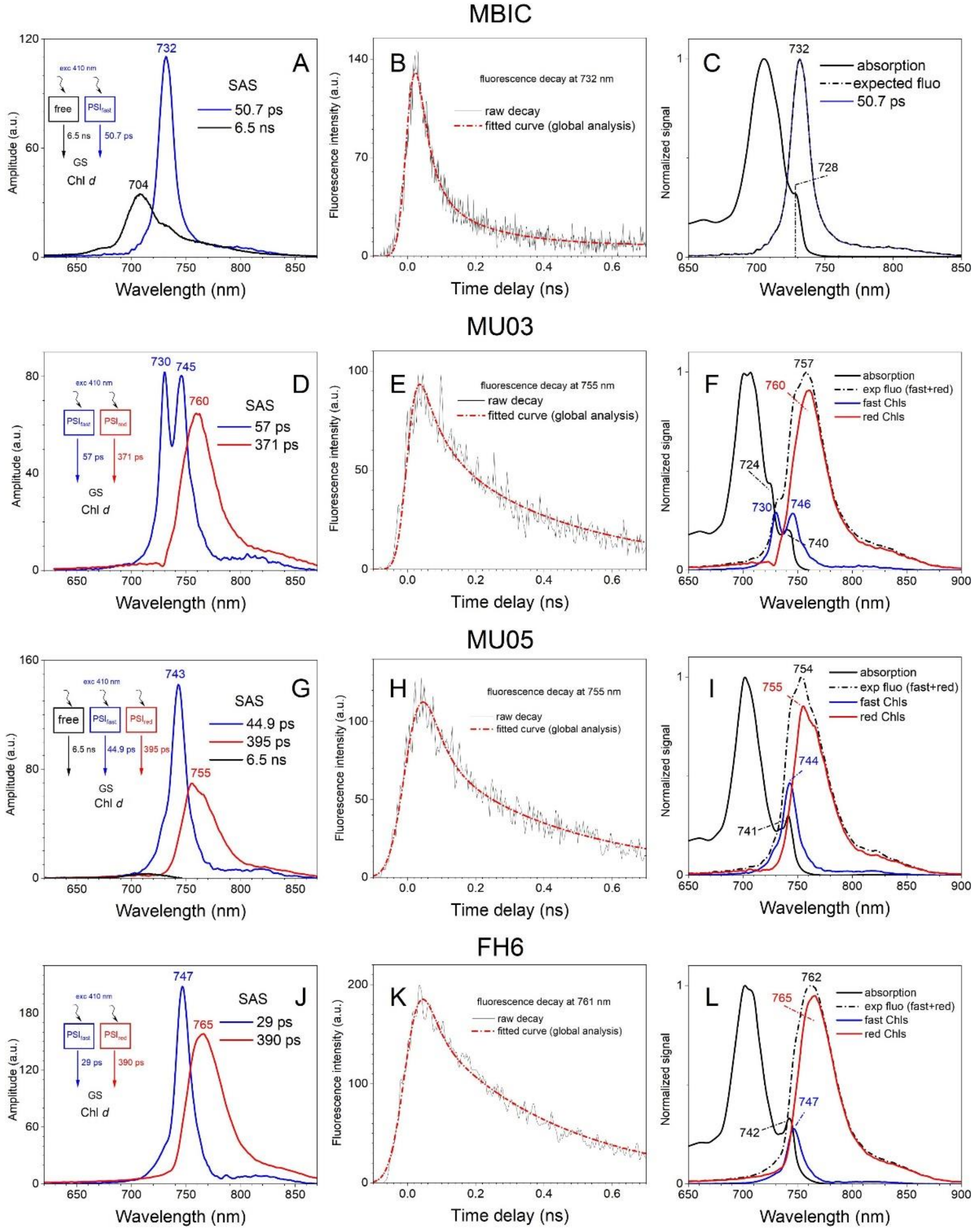

**Fig. S6.** Global analysis of 77 K TRF of spectral variants of PSI. (**A**, **D**, **G**, **J**) Global analysis of TRF map recorded at 77 K after excitation with laser pulse at 410 nm. The fitting models are indicated in the inserts. (**B**, **E**, **H**, **K**) TRF traces recorded at selected wavelength, accompanied by fits from global fitting. (**C**, **F**, **I**, **L**) Comparison of steady-state absorption and estimated steady-state fluorescence emission spectra (fluorescence from free chlorophylls omitted). Contributions of fluorescence from "fast" and "red" chlorophyll *d* molecules in the overall spectrum are provided. SAS – species associated spectra, GS – ground state.

**A**

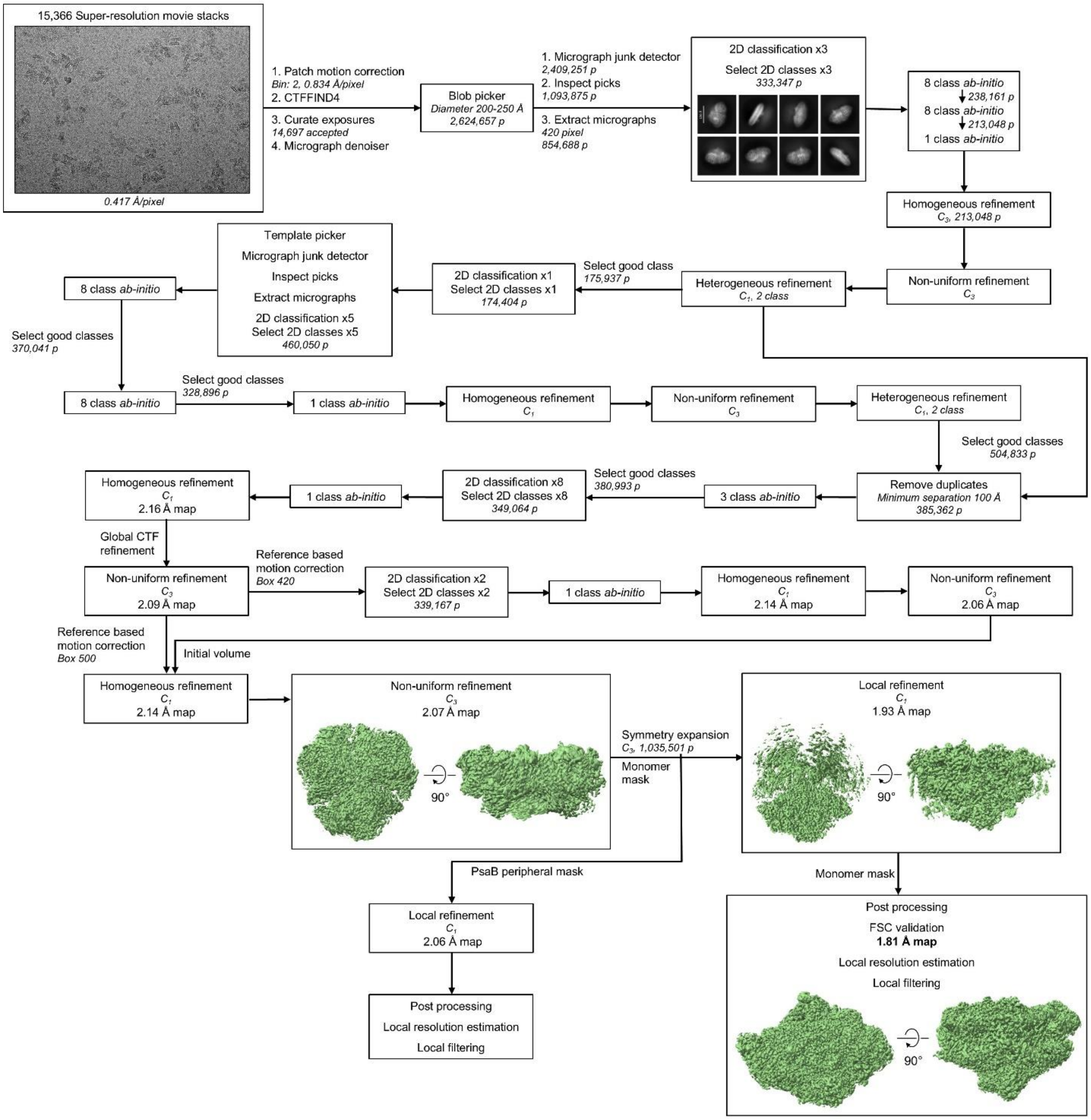
15,366 Super-resolution movie stacks
0.417 Å/pixel
1. Patch motion correction
Bin: 2, 0.834 Å/pixel
2. CTFFIND4
3. Curate exposures
14,697 accepted
4. Micrograph denoiser
Blob picker
Diameter 200-250 Å
2,624,657 p
1. Micrograph junk detector
2,409,251 p
2. Inspect picks
1,093,875 p
3. Extract micrographs
420 pixel
854,688 p
2D classification x3
Select 2D classes x3
333,347 p
8 class ab-initio
238,161 p
8 class ab-initio
213,048 p
1 class ab-initio
Homogeneous refinement
C3, 213,048 p
Non-uniform refinement
C3
Heterogeneous refinement
C1, 2 class
Select good class
175,937 p
2D classification x1
Select 2D classes x1
174,404 p
Template picker
Micrograph junk detector
Inspect picks
Extract micrographs
2D classification x5
Select 2D classes x5
460,050 p
8 class ab-initio
Select good classes
370,041 p
8 class ab-initio
Select good classes
328,896 p
1 class ab-initio
Homogeneous refinement
C1
Non-uniform refinement
C3
Heterogeneous refinement
C1, 2 class
Select good classes
504,833 p
Remove duplicates
Minimum separation 100 Å
385,362 p
3 class ab-initio
Select good classes
380,993 p
2D classification x8
Select 2D classes x8
349,064 p
1 class ab-initio
Homogeneous refinement
C1
2.16 Å map
Global CTF refinement
Non-uniform refinement
C3
2.09 Å map
Reference based motion correction
Box 420
2D classification x2
Select 2D classes x2
339,167 p
1 class ab-initio
Homogeneous refinement
C1
2.14 Å map
Non-uniform refinement
C3
2.06 Å map
Reference based motion correction
Box 500
Initial volume
Homogeneous refinement
C1
2.14 Å map
Non-uniform refinement
C3
2.07 Å map
90°
Symmetry expansion
C3, 1,035,501 p
Monomer mask
Local refinement
C1
1.93 Å map
90°
PsaB peripheral mask
Local refinement
C1
2.06 Å map
Post processing
Local resolution estimation
Local filtering
Monomer mask
Post processing
FSC validation
1.81 Å map
Local resolution estimation
Local filtering
90°

B

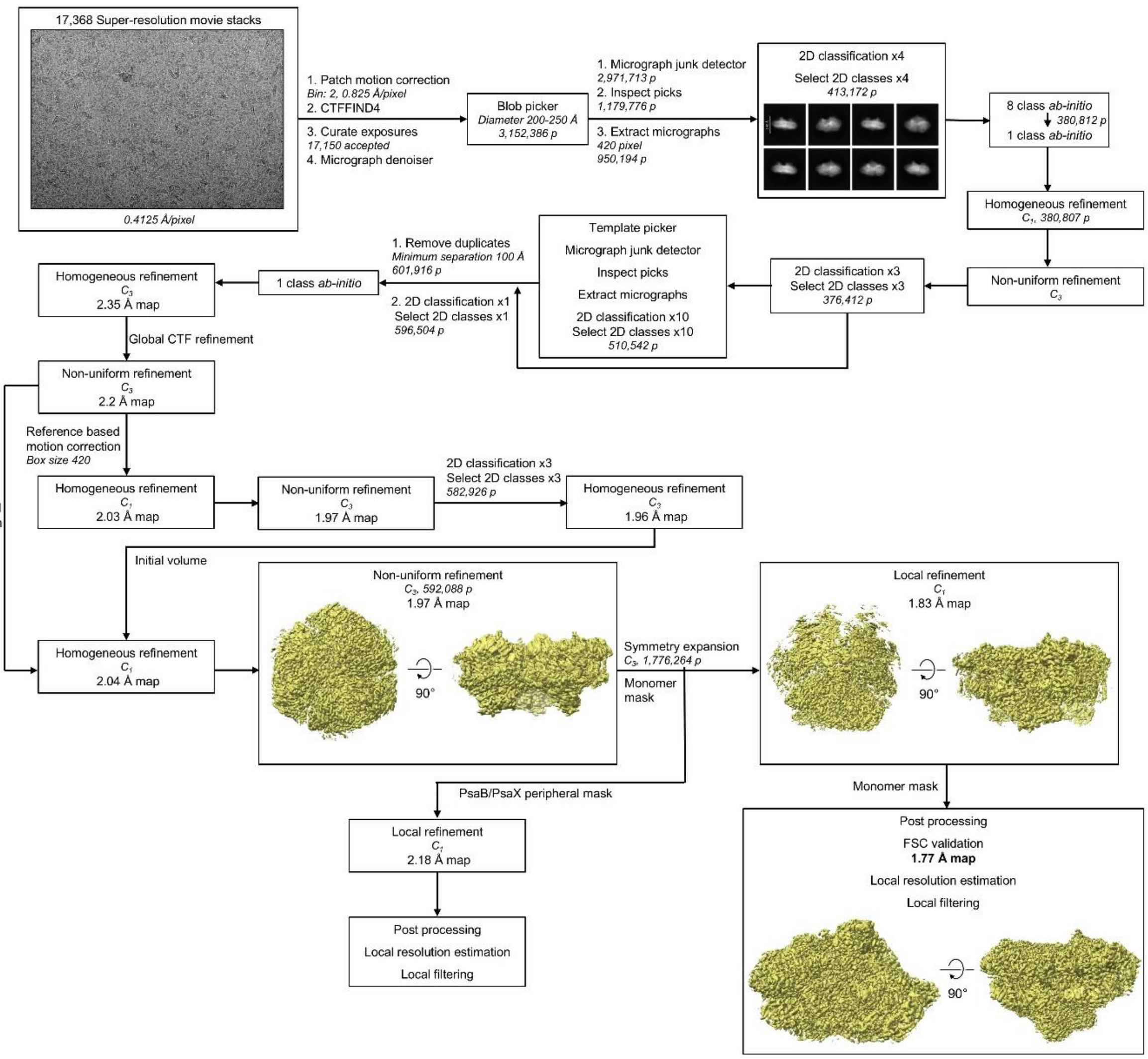

**Fig. S7.** Workflow of cryo-EM data processing in CryoSPARC (*23*). (**A**) MU03. (**B**) MU05.

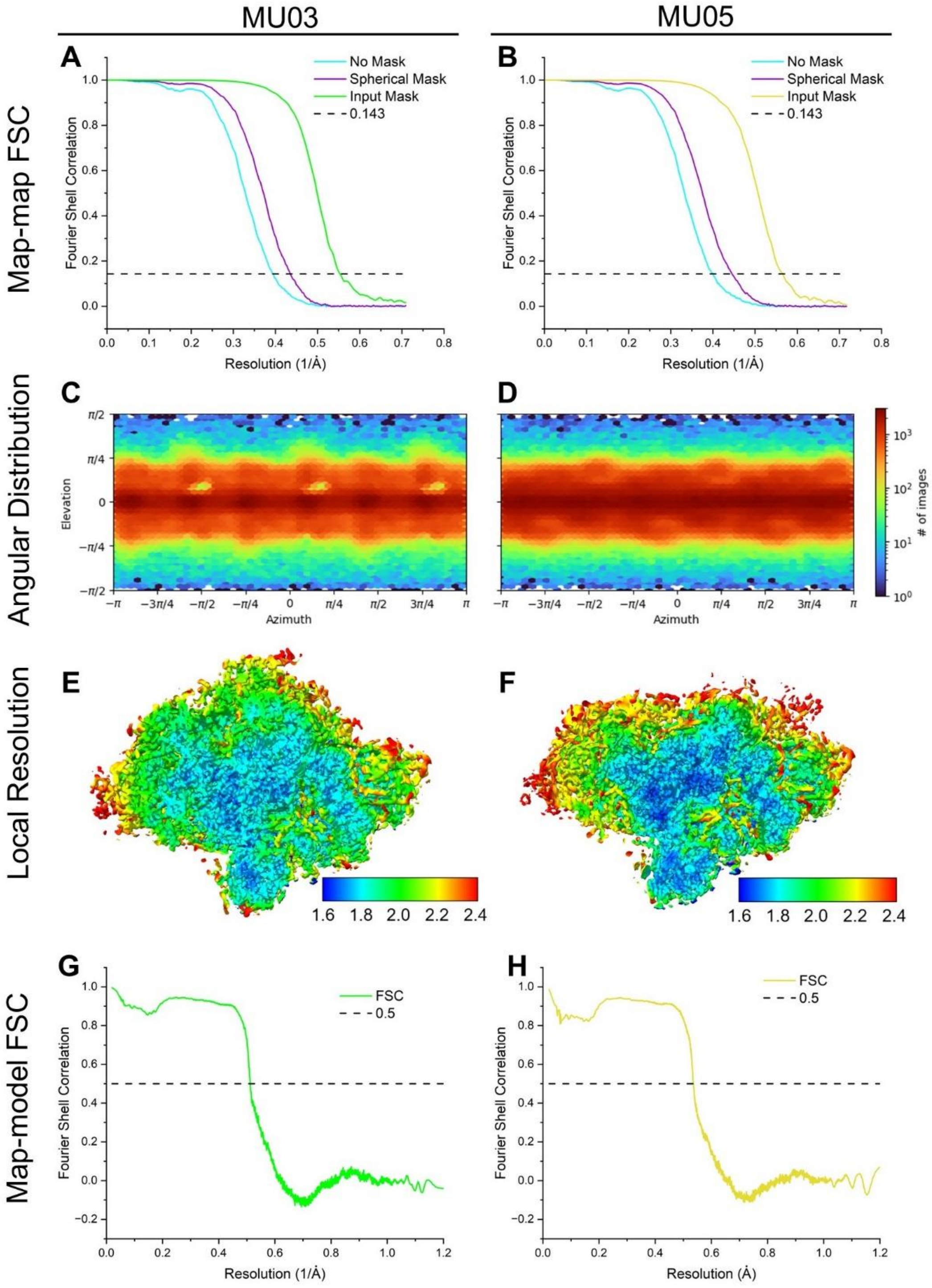

**Fig. S8.** Resolutions of cryo-EM data sets. (**A** and **B**) Gold-Standard Fourier Shell Correlation of final cryo-EM maps prior to local filtering. (**C** and **D**) Angular view distribution. (**E** and **F**) Slice through the locally filtered map colored by local resolution. The scale bar shows the local resolution in units of Å. (**G** and **H**) Map-model Fourier Shell Correlations.

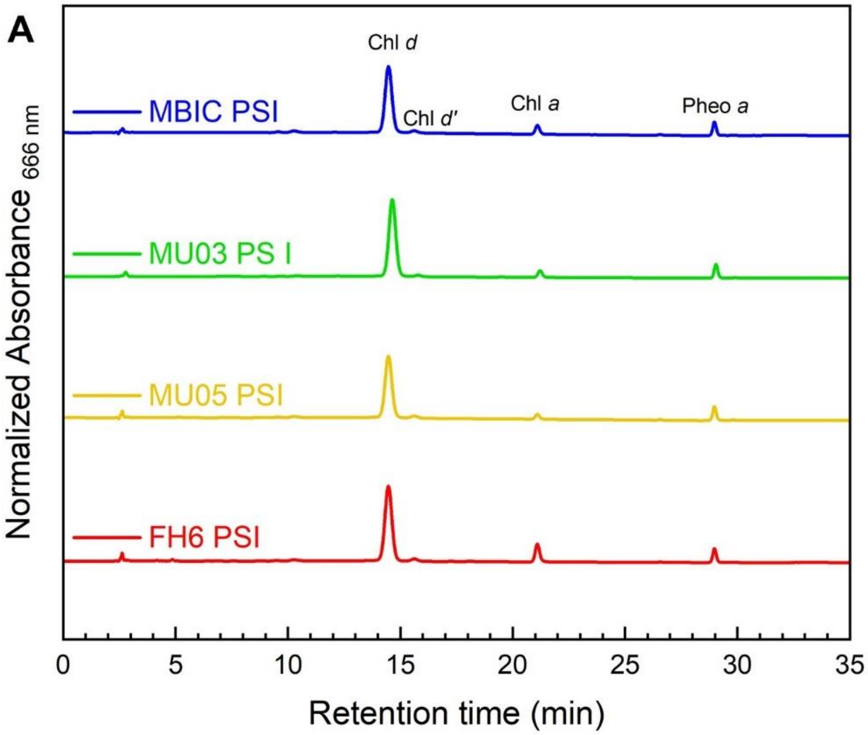

**B**

| | % Component | | | |
|---|---|---|---|---|
| | MBIC | MU03 | MU05 | FH6 |
| | tetrapyrroles | | | |
| **chlorophyll *d*** | 93.6 ± 0.80 | 95.0 ± 0.04 | 93.7 ± 0.40 | 92.2 ± 0.40 |
| **chlorophyll *d'*** | 2.16 ± 0.25 | 1.60 ± 0.00 | 2.60 ± 0.30 | 2.53 ± 0.05 |
| **chlorophyll *a*** | 1.68 ± 0.07 | 1.08 ± 0.02 | 0.98 ± 0.05 | 2.94 ± 0.15 |
| **pheophytin *a*** | 2.50 ± 0.47 | 2.37 ± 0.03 | 2.70 ± 0.50 | 2.30 ± 0.20 |
| | carotenoids | | | |
| **α-carotene** | 94.0 ± 0.30 | 91.2 ± 0.30 | 96.4 ± 0.70 | 94.3 ± 0.70 |
| **zeaxanthin** | 6.00 ± 0.30 | 8.80 ± 0.30 | 3.60 ± 0.60 | 5.70 ± 0.70 |

**Fig. S9.** HPLC pigment analysis of tetrapyrroles. (**A**) Chromatograms recorded at 666 nm of pigment extracts from PSI of MBIC, MU03, MU05, and FH6. All traces were normalized to the same pheophytin (Pheo) *a* content. (**B**) Pigment composition of *A. marina* PSI extracts. Data average of five (N=5) individual extraction except for MU03 (N=3).

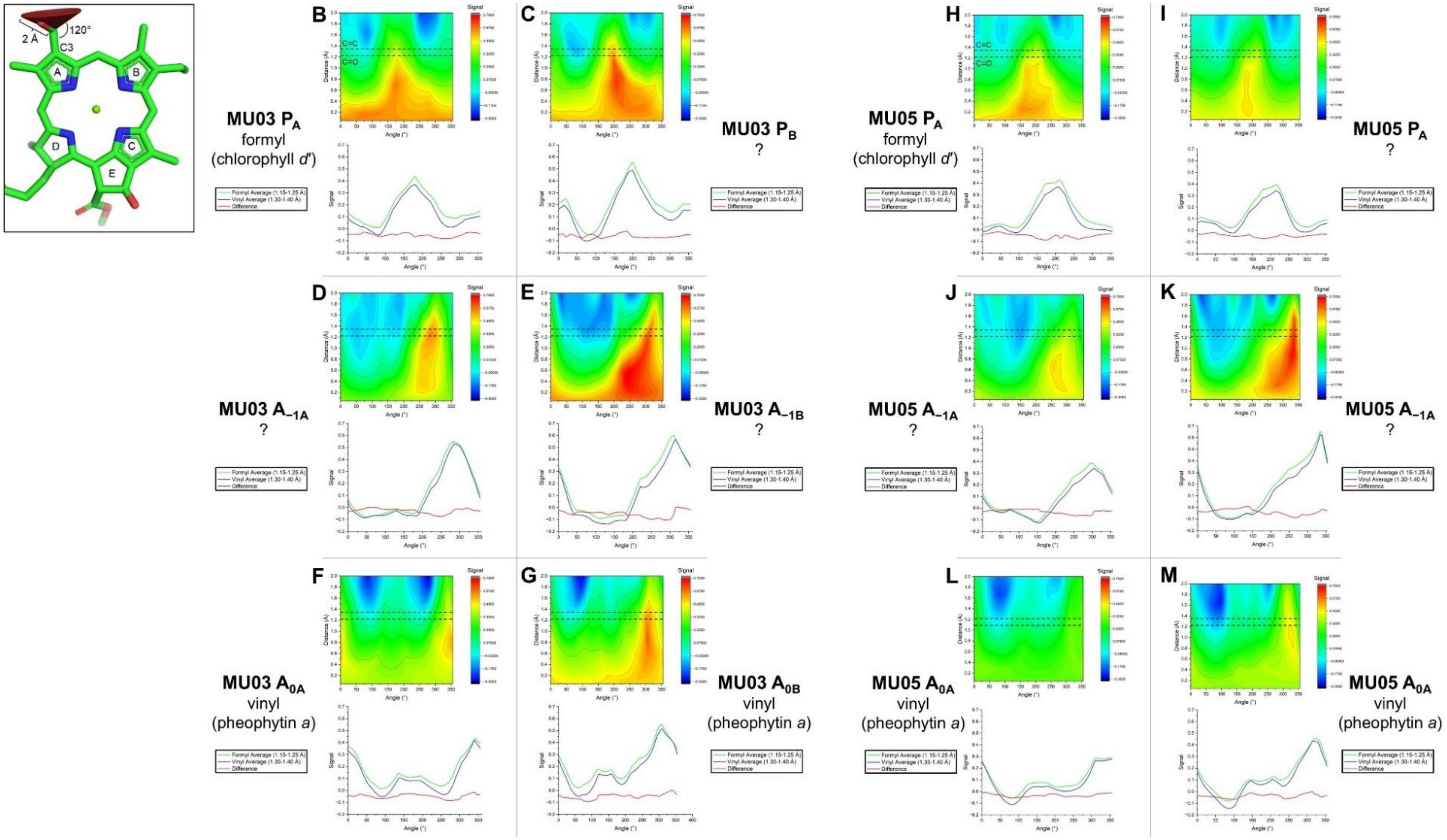

**Fig. S10.** Cone scan analysis of electron transfer chain tetrapyrroles. (**A**) A chlorophyll *d* molecule is shown with the cone probe atoms depicted as individual spheres at the C3 position. (**B-M**) The top panel shows the heat map of the cone scan. The X-axis is the angle from the tetrapyrrole ring where 0/360° is directed toward the pyrrole ring D and 180° is directed toward ring B. Horizontal lines show the expected C=O or C=C bind distance from the $C3_1$ atom. The bottom panel shows the average signal from 1.15-1.25 Å (green, approximate length of a C=O bond), the average signal from 1.30-1.40 Å (blue, approximate length of a C=C bond), and the subtraction of the former from the latter (red).

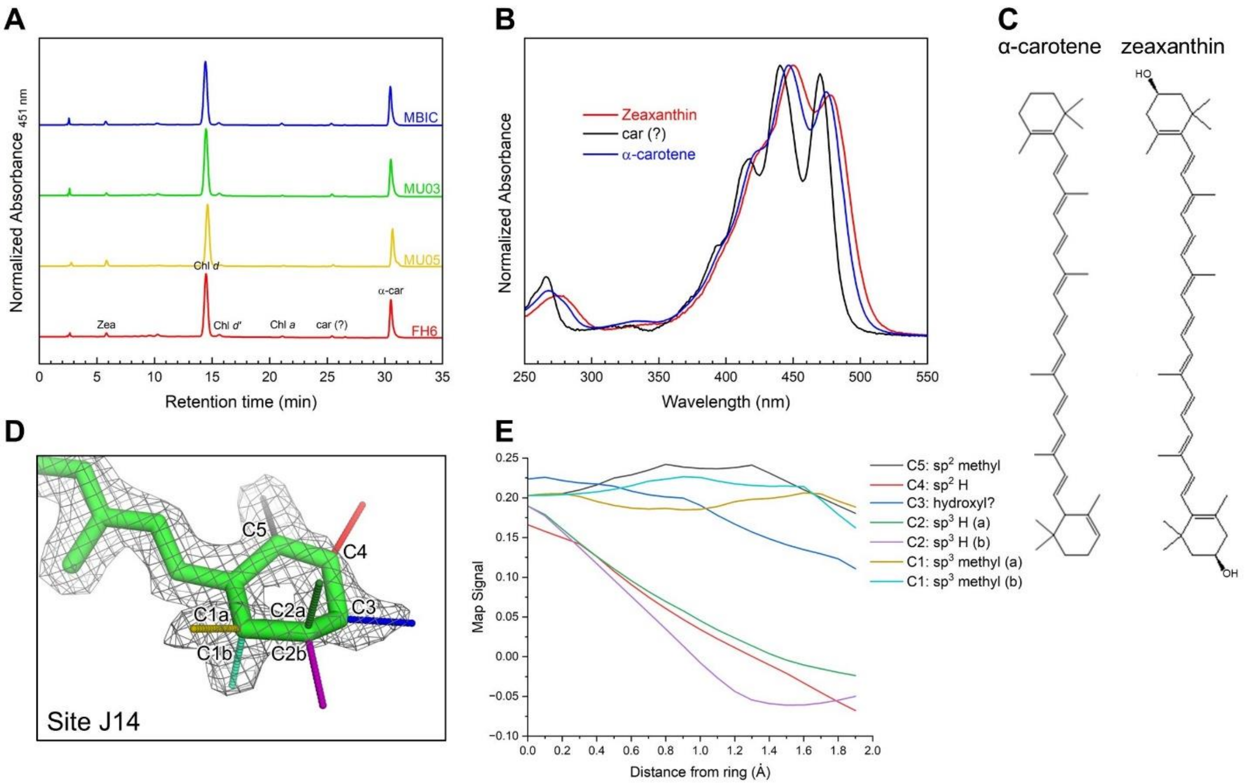

**Fig. S11.** Carotenoid analysis of MU03 and MU05 PSI. (**A**) HPLC chromatograms recorded at 451 nm of pigment extracts from PSI of MBIC, MU05, MU03, and FH6. Assignments of different molecules are shown on the FH6 PSI trace only. (**B**) Absorption spectra of the three carotenoids present in the pigment analysis normalized to the maximum absorbance. (**C**) Structures of α-carotene and zeaxanthin. (**D**) One ring of the carotenoid tentatively assigned as zeaxanthin in site J14 (naming based on Jordan et al. (*11*)) in MU03 PSI. (**E**) Map signal extending away from the ring shown in panel D.

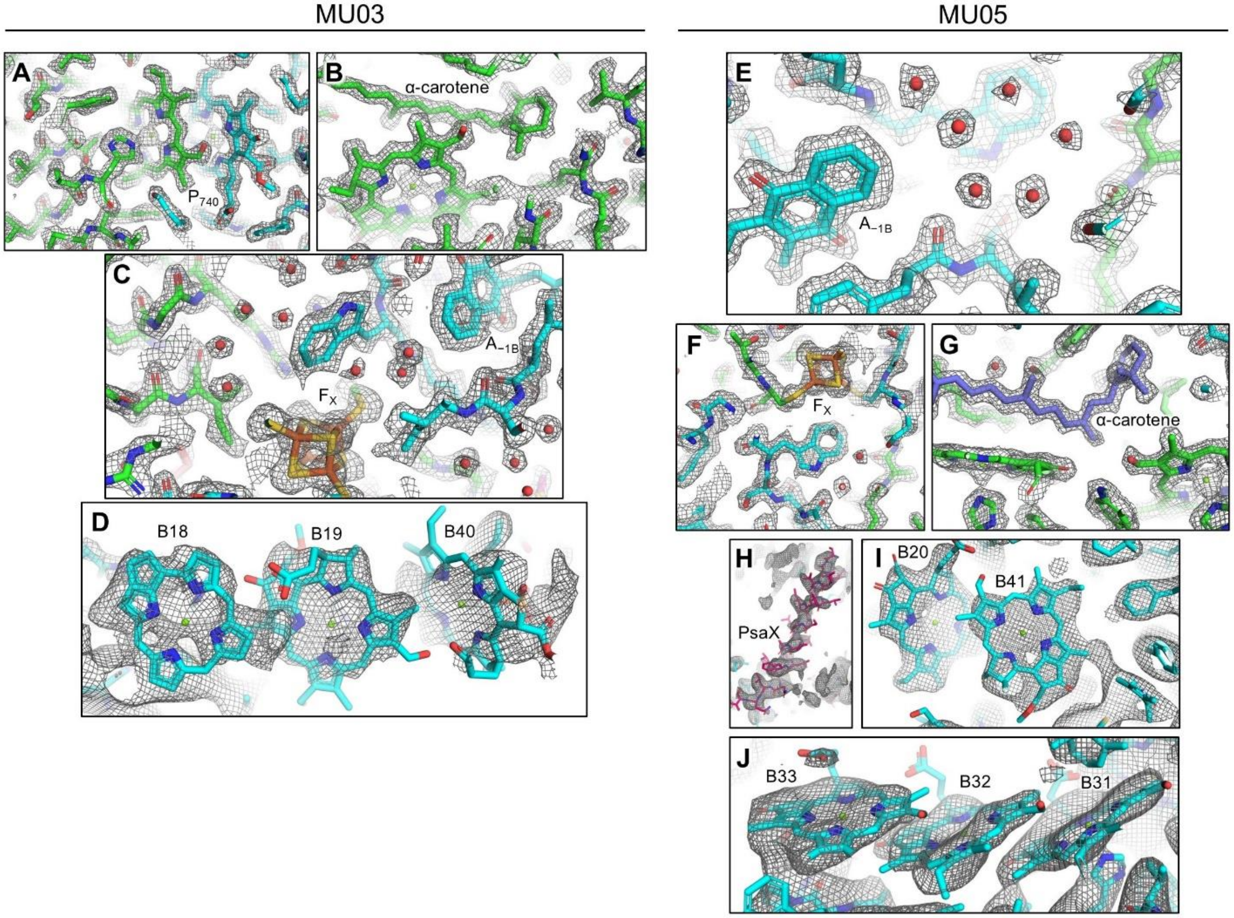

**Fig. S12.** Map examples for the MU03 and MU05 PSI cryo-EM structures. (**A**) Locally filtered MU03 map near $P_{740}$. (**B**) Locally filtered MU03 map near the headgroup of an α-carotene. (**C**) Locally filtered MU03 map near the $F_X/A_{-1B}$ region. (**D**) Unsharpened focused MU03 map near the three chlorophyll *d* molecules in sites B18, B19, and B40. (**E**) Locally filtered MU05 map near $A_{-1B}$. (**F**) Locally filtered MU05 map near $F_X$. (**G**) Locally filtered MU05 map near an α-carotene. (**H**) Unsharpened focused MU05 map PsaX. (**I**) Unsharpened focused MU05 map near the two chlorophyll *d* molecules in sites B20 and B41. (**J**) Unsharpened focused MU05 map near the three chlorophyll *d* molecules in sites B33, B32, and B31.

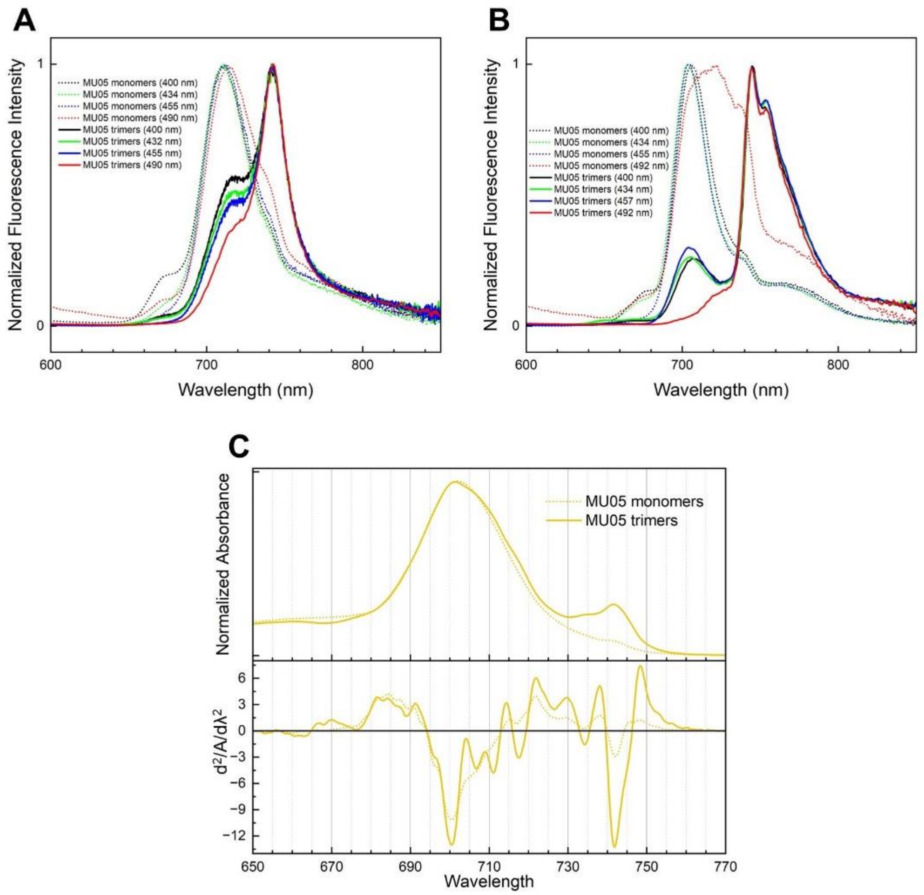

**Fig. S13.** Steady state spectroscopy comparing MU05 PSI trimers and monomers. (**A**) Room temperature fluorescence at different excitation wavelengths. (**B**) 77 K fluorescence at different excitation wavelengths. (**C**) Low temperature absorption spectrum and corresponding second derivative analysis. Samples used for fluorescence spectroscopy contain 60% glycerol.

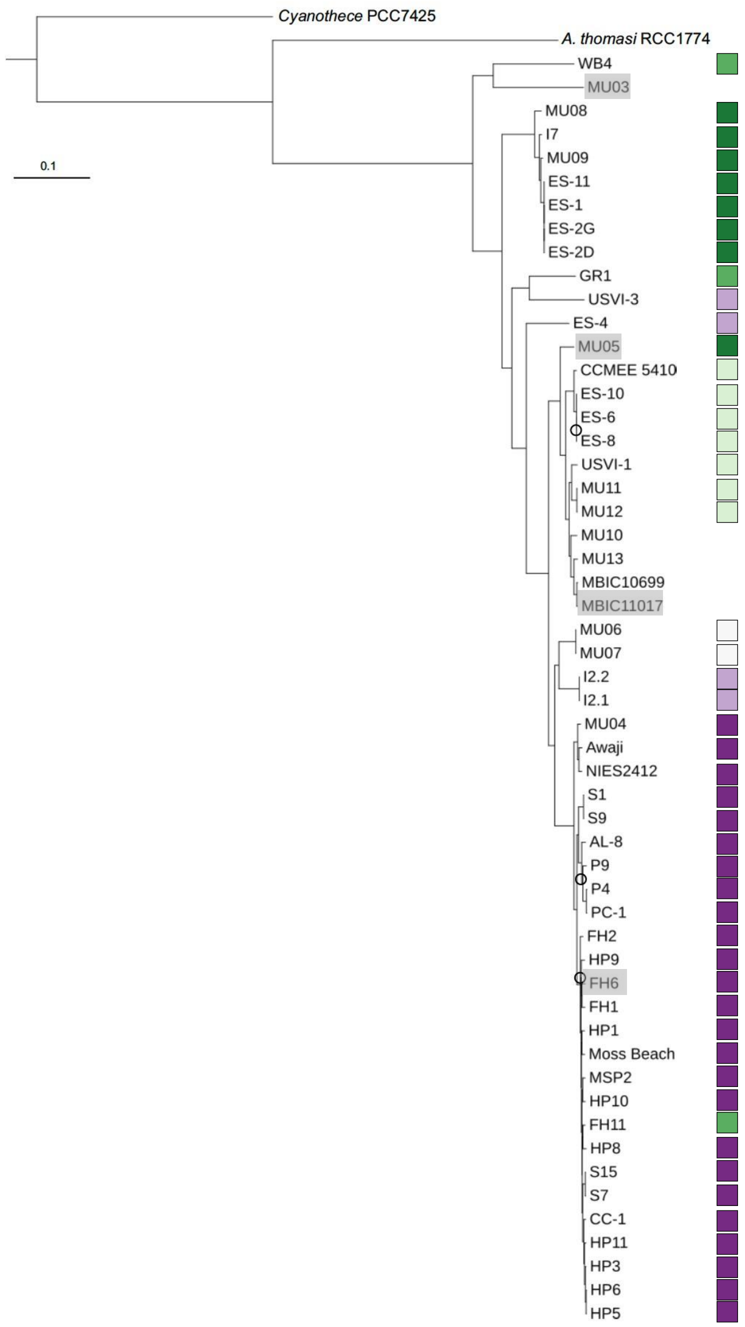

**Fig. S14.** Genome-wide amino acid phylogeny of *A. marina*. Strains investigated in this study are indicated by gray boxes. PsaX allele identities are mapped on the tree using the color coding in **Fig. 4**. The tree was reconstructed by maximum likelihood for a concatenation of 1,283 protein sequences from single-copy orthologs according to the JTT+F+R10 model of sequence evolution and outgroup-rooted with *Cyanothece* sp. PCC 7425. *Acaryochloris thomasi* RCC1774 produces chlorophyll *b* but not chlorophyll *d* and is sister to *A. marina*. Bootstrap support values for 1,000 ultrafast bootstrap replicates are >99% for all nodes, excluding those denoted by circles (80-99% support). Branch lengths are in units of expected number of amino acid substitutions per site. The strains we investigated herein are highlighted in grey.

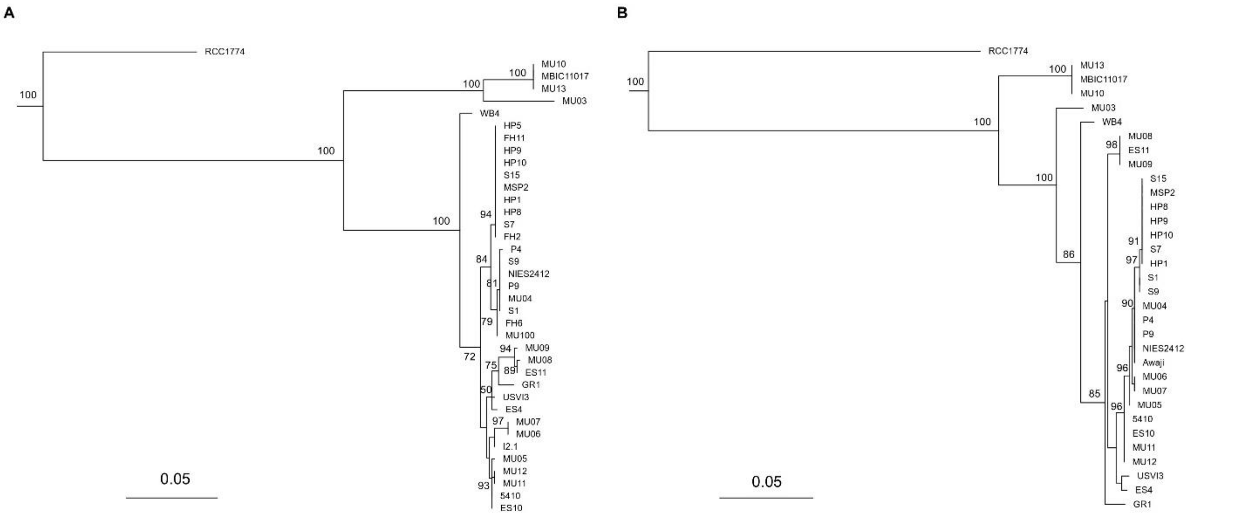

**Fig. S15.** Maximum likelihood tree for PsaA and PsaB from *A. marina.* (**A**) Maximum likelihood tree for *Acaryochloris* PsaB reconstructed with the MTZOA+F+I+R2 model of protein evolution and outgroup-rooted with sequences for *Thermosynechococcus vestitus* strain BP-1 and *Synechocystis* sp. PCC 6803. Ultrafast bootstrap values greater than 50% are shown at nodes. (**B**) Maximum likelihood tree for *Acaryochloris* PsaA reconstructed with the MTZOA+F+G4 model of protein evolution and outgroup-rooted with sequences for *Thermosynechococcus vestitus* strain BP-1 and *Synechocystis* sp. PCC 6803. Ultrafast bootstrap values greater than 50% are shown at nodes. The scale bar in both panels is in units of expected amino acid substitutions per amino acid site.

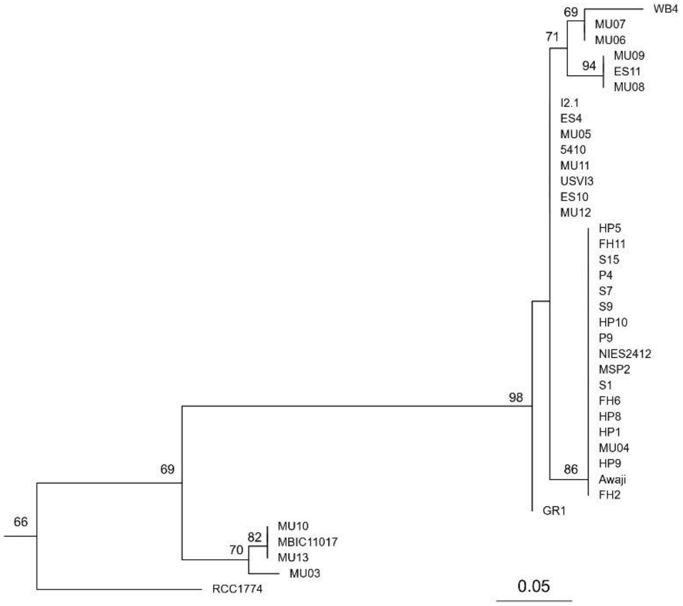

**Fig. S16.** Maximum likelihood tree for *Acaryochloris* PsaB amino acid positions 250-350 reconstructed with the Q.PFAM + R2 model of protein evolution and outgroup-rooted with sequences for *Thermosynechococcus vestitus* strain BP-1 and *Synechocystis* sp. PCC 6803. Ultrafast bootstrap values greater than 50% are shown at nodes. The scale bar is in units of expected amino acid substitutions per amino acid site.

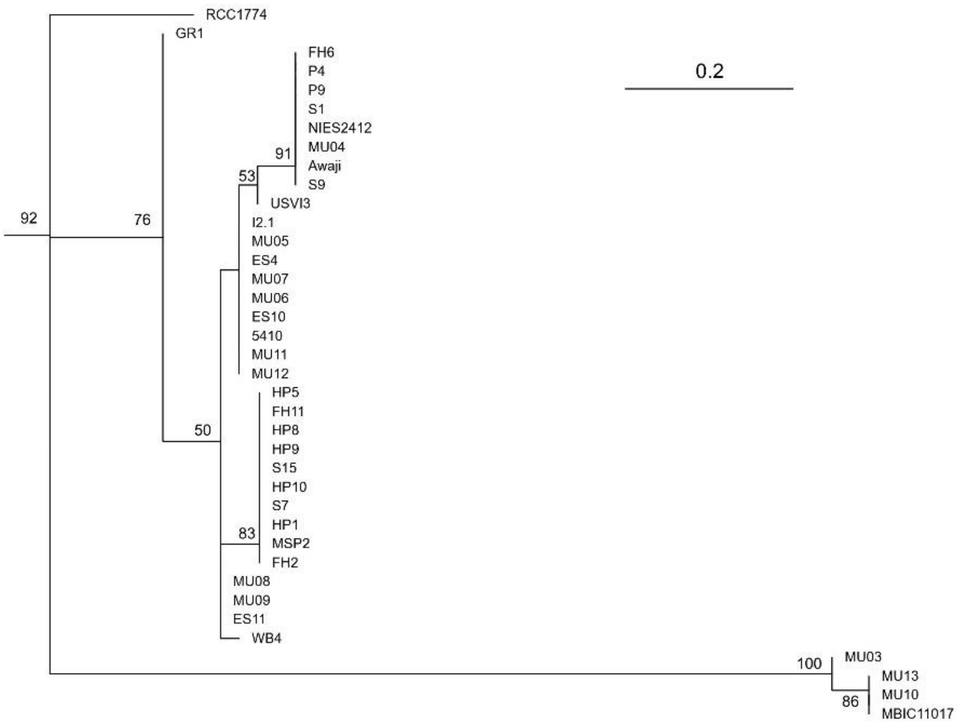

**Fig. S17.** Maximum likelihood tree for *Acaryochloris* PsaB amino acid positions 450-530 reconstructed with the WAG + G4 model of protein evolution and outgroup-rooted with sequences for *Thermosynechococcus vestitus* strain BP-1 and *Synechocystis* sp. PCC 6803. Ultrafast bootstrap values greater than 50% are shown at nodes. The scale bar is in units of expected amino acid substitutions per amino acid site.

**Table S1.** Mass spectrometry analysis of non-cross-linked PSI preparations from MBIC, MU05, MU03, and FH6. PSI subunits are highlighted in olive, PSII subunits are highlighted in beige, phycobiliproteins are highlighted in blue, and antenna proteins are highlighted in yellow. The spectral count is colored by a green (high) to white (low) gradient scheme.

**MBIC**

| Identified Proteins | Count |
|---|---|
| PsaD | 271 |
| PsaA | 246 |
| PsaE | 221 |
| PsaB | 196 |
| PsaF | 108 |
| PsaC | 104 |
| PsaL | 75 |
| PC | 56 |
| PsbB | 50 |
| PC | 41 |
| PC | 37 |
| PC | 36 |
| Linker | 33 |
| PsaC | 23 |
| PsbA | 14 |
| Linker | 11 |
| PsaD | 11 |
| Pcb-like | 11 |
| Pcb-like | 10 |
| Pcb-like | 7 |
| Ycf48 | 3 |
| Pcb-like | 3 |
| Pcb-like | 2 |
| Linker | 0 |
| PsbO | 0 |
| PsbQ | 0 |
| PsbP | 0 |
| PsbV | 0 |

**MU03**

| Identified Proteins | Count |
|---|---|
| PsaA | 490 |
| PsaD | 477 |
| PsaB | 455 |
| PsaE | 339 |
| CP47 | 223 |
| PsaC | 205 |
| PsaL | 175 |
| PsaF | 154 |
| CP43 | 91 |
| PsbA | 37 |
| PsbD | 30 |
| IsiA | 23 |
| Pcb-like | 21 |
| Pcb-like | 20 |
| Pcb-like | 20 |
| PcB-like | 16 |
| PsbP | 9 |
| PsbE | 8 |
| Psb27 | 8 |
| PsbW | 7 |
| Pcb-like | 5 |

**MU05**

| Protein | Count |
|---|---|
| PsaA | 303 |
| PsaD | 206 |
| PsaB | 152 |
| PsaC | 126 |
| PsbB | 93 |
| PsaF | 82 |
| PsaL | 76 |
| PsbC | 53 |
| Pcb-like | 47 |
| Pcb-like | 40 |
| Pcb-like | 32 |
| PsaE | 21 |
| PsbA | 20 |
| Pcb-like | 14 |
| PsbD | 12 |
| PsbD | 12 |
| Pcb-like | 12 |
| Pcb-like | 12 |
| Pcb-like | 12 |
| PsbD | 11 |
| Pcb-like | 9 |
| IsiA | 9 |
| Pcb-like | 8 |
| PsbE | 5 |
| PsbF | 4 |
| Psb27 | 4 |
| PsbP | 2 |
| PsbD | 0 |
| PsbU | 0 |

**FH6**

| Identified Proteins | Count |
|---|---|
| PsaA | 299 |
| PsaD | 185 |
| PsaB | 168 |
| PsaF | 164 |
| PsaE | 107 |
| PsaC | 105 |
| PsaB | 99 |
| PsaL | 56 |
| PsbC | 40 |
| Pcb-like | 29 |
| Pcb-like | 25 |
| PsbD | 22 |
| PsbA2 | 15 |
| PsbP | 7 |
| PsbE2 | 4 |
| PsaK2 | 4 |
| Psb27 | 2 |
| PsbW | 0 |
| PsbH | 0 |
| PsbV2 | 0 |

**Table S2.** Sequence identity matrix of PSII core subunits.

**PsbA (D1)**

| | MBIC PsbA1 | MU05 PsbA1 | FH6 PsbA | MU03 PsbA | MU05 PsbA3 | FH6 PsbA3 | MBIC PsbA2 | MU05 PsbA2 | FH6 PsbA2 |
|---|---|---|---|---|---|---|---|---|---|
| MBIC PsbA1 | | 100 | 100 | 99 | 62 | 62 | 62 | 62 | 62 |
| MU05 PsbA1 | 100 | | 100 | 99 | 62 | 62 | 62 | 62 | 62 |
| FH6 PsbA | 100 | 100 | | 99 | 62 | 62 | 62 | 62 | 62 |
| MU03 PsbA | 99 | 99 | 99 | | 62 | 62 | 62 | 62 | 62 |
| MU05 PsbA3 | 62 | 62 | 62 | 62 | | 100 | 81 | 84 | 84 |
| FH6 PsbA3 | 62 | 62 | 62 | 62 | 100 | | 81 | 84 | 84 |
| MBIC PsbA2 | 62 | 62 | 62 | 62 | 81 | 81 | | 88 | 88 |
| MU05 PsbA2 | 62 | 62 | 62 | 62 | 84 | 84 | 88 | | 100 |
| FH6 PsbA2 | 62 | 62 | 62 | 62 | 84 | 84 | 88 | 100 | |

**PsbB (CP47)**

| | MBIC PsbB | MU05 PsbB | MU03 PsbB | FH6 PsbB |
|---|---|---|---|---|
| MBIC PsbB | | 98 | 99 | 97 |
| MU05 PsbB | 98 | | 98 | 99 |
| MU03 PsbB | 99 | 98 | | 98 |
| FH6 PsbB | 97 | 99 | 98 | |

**PsbC (CP43)**

| | MBIC PsbC | MU05 PsbC | MU03 PsbC | FH6 PsbC |
|---|---|---|---|---|
| MBIC PsbC | | 99 | 96 | 99 |
| MU05 PsbC | 99 | | 96 | 99 |
| MU03 PsbC | 96 | 96 | | 96 |
| FH6 PsbC | 99 | 99 | 96 | |

**PsbD (D2)**

| | MBIC PsbD2 | MU03 PsbD1 | MU03 PsbD2 | MBIC PsbD1 | MU05 PsbD | FH6 PsbD |
|---|---|---|---|---|---|---|
| MBIC PsbD2 | | 94 | 94 | 94 | 93 | 93 |
| MU03 PsbD1 | 94 | | 100 | 98 | 99 | 99 |
| MU03 PsbD2 | 94 | 100 | | 98 | 99 | 99 |
| MBIC PsbD1 | 94 | 98 | 98 | | 99 | 99 |
| MU05 PsbD | 93 | 99 | 99 | 99 | | 100 |
| FH6 PsbD | 93 | 99 | 99 | 99 | 100 | |

**Table S3.** Sequence identity matrix of PSI subunits.

**PsaA**

| | MBIC PsaA | MU03 PsaA | MU05 PsaA | FH6 PsaA |
|---|---|---|---|---|
| MBIC PsaA | | 93 | 93 | 93 |
| MU03 PsaA | 93 | | 96 | 96 |
| MU05 PsaA | 93 | 96 | | 99 |
| FH6 PsaA | 93 | 96 | 99 | |

**PsaB**

| | MBIC PsaB | MU03 PsaB | MU05 PsaB | FH6 PsaB |
|---|---|---|---|---|
| MBIC PsaB | | 94 | 90 | 89 |
| MU03 PsaB | 94 | | 88 | 88 |
| MU05 PsaB | 90 | 88 | | 99 |
| FH6 PsaB | 89 | 88 | 99 | |

**PsaC**

| | MBIC PsaC | MU03 PsaC | MU05 PsaC | FH6 PsaC |
|---|---|---|---|---|
| MBIC PsaC | | 98 | 100 | 99 |
| MU03 PsaC | 98 | | 98 | 99 |
| MU05 PsaC | 100 | 98 | | 99 |
| FH6 PsaC | 99 | 99 | 99 | |

**PsaD**

| | MBIC PsaD | MU03 PsaD | MU05 PsaD | FH6 PsaD |
|---|---|---|---|---|
| MBIC PsaD | | 91 | 99 | 99 |
| MU03 PsaD | 91 | | 91 | 91 |
| MU05 PsaD | 99 | 91 | | 98 |
| FH6 PsaD | 99 | 91 | 98 | |

**PsaE**

| | MBIC PsaE | MU03 PsaE | MU05 PsaE | FH6 PsaE |
|---|---|---|---|---|
| MBIC PsaE | | 89 | 94 | 95 |
| MU03 PsaE | 89 | | 89 | 86 |
| MU05 PsaE | 94 | 89 | | 95 |
| FH6 PsaE | 95 | 86 | 95 | |

**PsaF**

| | MBIC PsaF | MU03 PsaF | MU05 PsaF | FH6 PsaF |
|---|---|---|---|---|
| MBIC PsaF | | 89 | 92 | 87 |
| MU03 PsaF | 89 | | 90 | 87 |
| MU05 PsaF | 92 | 90 | | 95 |
| FH6 PsaF | 87 | 87 | 95 | |

**PsaI**

| | MBIC PsaI | MU03 PsaI | MU05 PsaI | FH6 PsaI |
|---|---|---|---|---|
| MBIC PsaI | | 88 | 100 | 100 |
| MU03 PsaI | 88 | | 88 | 88 |
| MU05 PsaI | 100 | 88 | | 100 |
| FH6 PsaI | 100 | 88 | 100 | |

**PsaJ**

| | MBIC PsaJ | MU03 PsaJ | MU05 PsaJ | FH6 PsaJ |
|---|---|---|---|---|
| MBIC PsaJ | | 89 | 95 | 97 |
| MU03 PsaJ | 89 | | 95 | 92 |
| MU05 PsaJ | 95 | 95 | | 97 |
| FH6 PsaJ | 97 | 92 | 97 | |

**PsaK**

| | MBIC PsaK1 | MU03 PsaK1 | MU05 PsaK1 | FH6 PsaK1 | MBIC PsaK2 | MU03 PsaK2 | MU05 PsaK2 | FH6 PsaK2 |
|---|---|---|---|---|---|---|---|---|

| | | | | | | | | |
|---|---|---|---|---|---|---|---|---|
| MBIC PsaK1 | | 87 | 98 | 97 | 35 | 41 | 33 | 35 |
| MU03 PsaK1 | 87 | | 86 | 87 | 35 | 37 | 33 | 35 |
| MU05 PsaK1 | 98 | 86 | | 98 | 40 | 41 | 35 | 40 |
| FH6 PsaK1 | 97 | 87 | 98 | | 38 | 40 | 37 | 38 |
| MBIC PsaK2 | 35 | 35 | 40 | 38 | | 85 | 92 | 99 |
| MU03 PsaK2 | 41 | 37 | 41 | 40 | 85 | | 83 | 85 |
| MU05 PsaK2 | 33 | 33 | 35 | 37 | 92 | 83 | | 91 |
| FH6 PsaK2 | 35 | 35 | 40 | 38 | 99 | 85 | 91 | |

**PsaL**

| | MBIC PsaL | MU03 PsaL | MU05 PsaL | FH6 PsaL |
|---|---|---|---|---|
| MBIC PsaL | | 90 | 95 | 89 |
| MU03 PsaL | 90 | | 91 | 95 |
| MU05 PsaL | 95 | 91 | | 91 |
| FH6 PsaL | 89 | 95 | 91 | |

**PsaM**

| | MBIC PsaM | MU03 PsaM | MU05 PsaM | FH6 PsaM |
|---|---|---|---|---|
| MBIC PsaM | | 97 | 100 | 97 |
| MU03 PsaM | 97 | | 97 | 94 |
| MU05 PsaM | 100 | 97 | | 97 |
| FH6 PsaM | 97 | 94 | 97 | |

**PsaX**

| | MU05 PsaX | FH6 PsaX |
|---|---|---|
| MU05 PsaX | | 50 |
| FH6 PsaX | 50 | |

**Table S4.** Assigned Stokes shifts based on spectroscopic data.

| | Absorption maximum (nm) | Fluorescence maximum (nm) | Stokes shift (nm) |
|---|---|---|---|
| MBIC | 729 | 732 | 3 |
| MU03 | 726 | 730 | 4 |
| | 740 | 745 | 5 |
| | 745 | 763 | 18 |
| MU05 | 742 | 745 | 3 |
| | 745 | 754 | 9 |
| FH6 | 742 | 745 | 3 |
| | 745 | 763 | 18 |

**Table S5.** Cryo-EM data collection, refinement, and validation statistics. *Super-resolution pixel size. †C3 symmetry was imposed during refinement prior to C3 symmetry expansion. Subsequent refinement was performed without symmetry. ‡Based on refinement in the maps generated from local filtering. °Prior to local filtering.

| | MU03 PSI | MU05 PSI |
|---|---|---|
| **Data Collection and Processing** | | |
| Magnification | ×105,000 | ×105,000 |
| Voltage (kV) | 300 | 300 |
| Electron Exposure ($e^{-}/Å^2$) | 50 | 50 |
| Defocus Range (μm) | −0.7 to −1.9 | −0.7 to −1.9 |
| Pixel Size (Å)* | 0.417 | 0.413 |
| Symmetry Imposed† | C3 | C3 |
| Initial Particle Images (no.) | 2,624,657 | 3,152,386 |
| Final Particle Images (no.) | 1,035,501 | 1,776,264 |
| Map Resolution (Å) | 1.81 | 1.77 |
| FSC Threshold | 0.143 | 0.143 |
| **Refinement‡** | | |
| Initial Model Used (PDB) | 7COY | 7COY |
| Model Resolution (Å)‡ | 1.95 | 1.87 |
| FSC Threshold | 0.5 | 0.5 |
| Map Resolution Range (Å, minimum to $75^{th}$ percentile) | 1.65-2.21 | 1.56-2.36 |
| Map-sharpening *B* Factor ($Å^2$)° | −57.5 | −48.2 |
| Model Composition | | |
| Non-hydrogen Atoms | 24,488 | 24,061 |
| Protein Residues | 2,190 | 2,159 |
| Ligands | 132 | 128 |
| B Factors Mean ($Å^2$) | | |
| Protein | 36.09 | 37.57 |
| Ligands | 39.29 | 42.72 |
| R.M.S. Deviations | | |
| Bond Lengths (Å) | 0.028 | 0.026 |
| Bond Angles (°) | 2.071 | 2.168 |
| **Validation** | | |
| MolProbity | 1.30 | 1.54 |
| Clashscore | 4.17 | 6.67 |
| Rotamer Outliers (%) | 0.11 | 0.29 |
| Ramachandran Plot | | |
| Favored (%) | 97.50 | 97.05 |
| Allowed (%) | 2.41 | 2.81 |
| Disallowed (%) | 0.09 | 0.14 |

**Table S6.** Cofactors modeled in each monomer of the *A. marina* PSI structures.

| Cofactor | PDB code | MU03 PSI | MU05 PSI |
|---|---|---|---|
| Chlorophyll *d* | CL7 | 91 | 91 |
| α-carotene | WVN | 17 | 17 |
| β-DDM | LMT | 8 | 6 |
| Phosphatidyl glycerol | LHG | 4 | 3 |
| [4Fe-4S] cluster | SF4 | 3 | 3 |
| Pheophytin *a* | PHO | 2 | 2 |
| Phylloquinone | PQN | 2 | 2 |
| Distearoyl-monogalactosyl-diglyceride | LMG | 2 | 1 |
| Calcium cation | CA | 1 | 2 |
| Chlorophyll *d′* | G9R | 1 | 1 |
| Zeaxanthin | K3I | 1 | 0 |

**Data S1** (separate files).
Förster-type excitation energy transfer calculations for PSI antenna chlorophylls. Includes pairwise coupling rate matrices (*_forster_rates.txt), orientation factor matrices (*_k_matrix.txt), and filtered high-coupling interaction tables used for visualization (*_fast_rates.txt) for each PSI structure analyzed.
